# SPtsAnalysis: a high-throughput super-resolution single particle trajectory analysis to reconstruct organelle dynamics and membrane re-organization

**DOI:** 10.1101/2021.09.18.460892

**Authors:** P. Parutto, J. Heck, M. Lu, C. Kaminski, M. Heine, E. Avezov, D. Holcman

## Abstract

Super-resolution imaging can generate thousands of single-particle trajectories. These data can potentially reconstruct subcellular organization and dynamics, as well as measure disease-linked changes. However, computational methods that can derive quantitative information from such massive datasets are currently lacking. Here we present data analysis and algorithms that are broadly applicable to reveal local binding and trafficking interactions and organization of dynamic sub-cellular sites. We applied this analysis to the endoplasmic reticulum and neuronal membrane. The method is based on spatio-temporal time window segmentation that explores data at multiple levels and detects the architecture and boundaries of high density regions in areas that are hundreds of nanometers. By statistical analysis of a large number of datapoints, the present method allows measurements of nano-region stability. By connecting highly dense regions, we reconstructed the network topology of the ER, as well as molecular flow redistribution, and the local space explored by trajectories. Segmenting trajectories at appropriate scales extracts confined trajectories, allowing quantification of dynamic interactions between lysosomes and the ER. A final step of the method reveals the motion of trajectories relative to the ensemble, allowing reconstruction of dynamics in normal ER and the atlastin-null mutant. Our approach allows users to track previously inaccessible large scale dynamics at high resolution from massive datasets. The SPtsAnalysis algorithm is available as an ImageJ plugin that can be applied by users to large datasets of overlapping trajectories and offer a standard of SPTs metrics.

## 1 Introduction

Sub-cellular compartments are focused sites where large numbers of molecules dynamically interact to support cellular function [1]. The trajectories of ions and proteins as they move between the cytoplasm, plasma membrane and organelles are critical to cellular function [2, 3]. These dynamics occur at the Endoplasmic Reticulum (ER) [4, 5, 6], the mitochondrial network, endosomes and lysosomes and microtubules, and are impacted by the local properties of these different environments [7]. Several experimental paradigms can measure these constitutive molecular motions at subcellular sites, including the reciprocal fluorescence recovery after photobleaching (FRAP) [8, 9] which locally depletes fluorescence and measures time scales and fraction of recovery. Analysis of these data can reveal trafficking at a population level. In contrast, photoactivation [10] consists of activating molecules in a local region of the cell and reveals their spread over a transient time-frame. Combined with diffusion modeling and stochastic simulations, various biophysical properties can be measured, including diffusion coefficients and the fraction and time scale of recovery [11]. These methods provide information on the dynamic function of organelles, but are insufficient to identify and reconstruct high density regions. These approaches also cannot examine phase separation stability or the local spaces is explored by molecules at a nanoscale resolution. Statistical analysis [12, 13] of a large ensemble of super-resolution single-particle trajectories SPTs (Fig. 1A-B), has the potential to reveal local molecular interactions. Molecules are not uniformly distributed inside a cell but instead form heterogeneous aggregates, possibly in phase-separated nanodomains, characterized by high density regions (HDRs). Such regions are characterized by reduced velocity movement of molecules and confinement. These local areas can be enriched with calcium channels at neuronal synapses, store-operated calcium entry receptors such as STIM1 on spine apparatus, and can also be found at ER nodes [14, 15, 16, 17]. Interestingly, these ubiquitous structures are transient, yet persist with a time scale that is longer than that associated with molecular trafficking. In short, many interactions that are critical to the cell are regulated and controlled by sub-cellular mechanisms that currently cannot be easily captured by quantitative analysis.

**Fig. 1.**
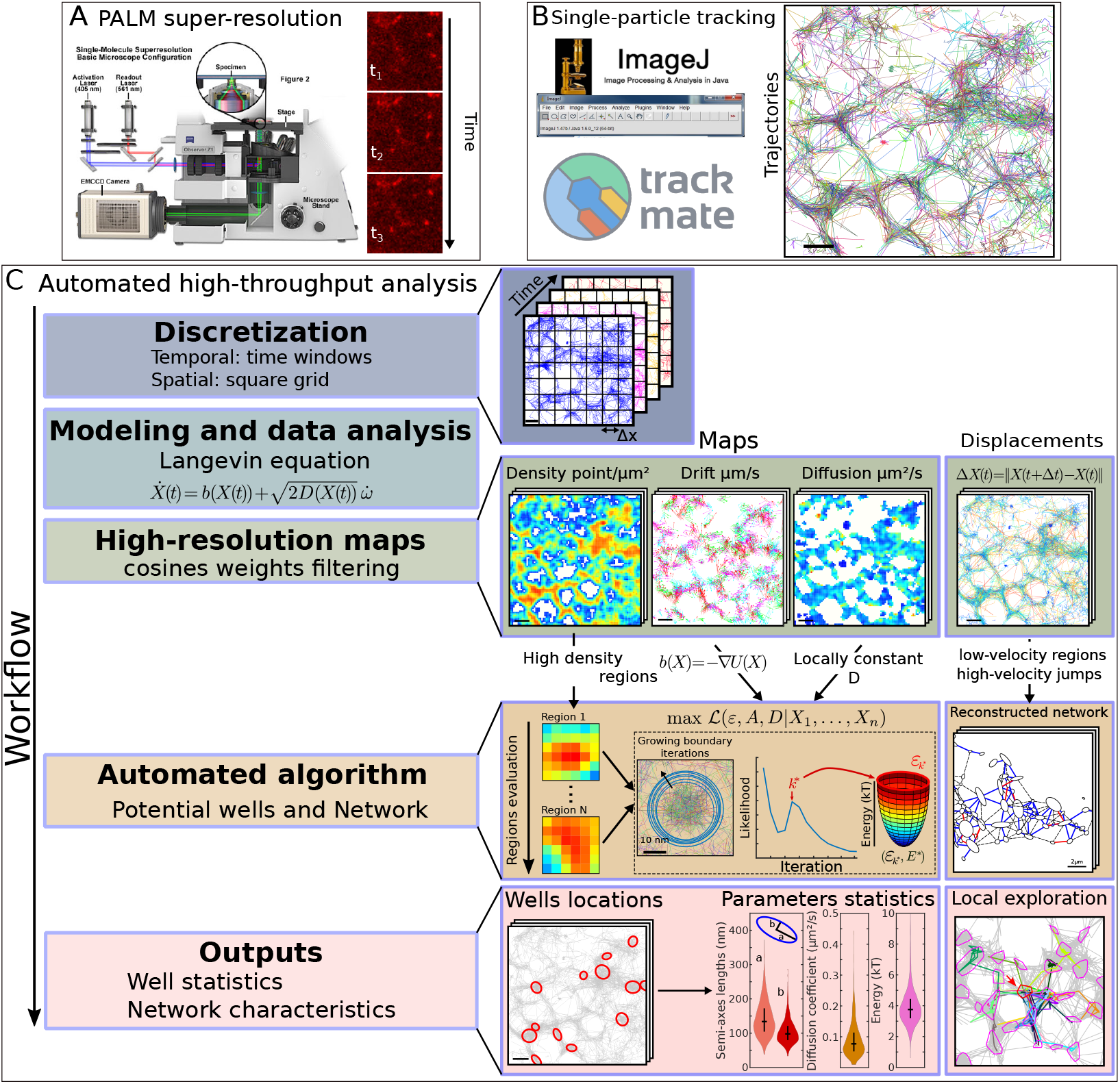
High-throughput SPT analysis pipeline. **High-throughput automated trajectory analysis workflow. A.** Acquisition device and raw data of a single-particle experiment. **B.** Raw data from A are transformed into trajectories using classical softwares such as Trackmate available as an ImageJ plugin. **C.** Schematic description of the high-throughput analysis implemented here: trajectories are first discretized spatially using a square grid and a time-windows generate a segmentation in time. Thet are interpreted using the Langevin equation, allowing us to generate high-resolution maps of the local trajectory motion. High-density / low-velocity regions of the maps are generated by automated machine-learning type algorithms to detect potential wells and to reconstruct network. Outputs consist in location maps of wells, their statistics and reconstructed network allowing to analyse how trajectories locally exploration the nanophysiology scale.

To determine the underlying physical properties of molecular trafficking, various computational modeling methods have been developed to analyze SPTs. These models include those based on classical free [18] and confined [19, 20, 9, 21] diffusion, active deterministic motion or a mixture of deterministic and stochastic models [22]. Based on this theoretical framework, analysis of SPTs has revealed the dynamics of local chromatin organization in the nucleus [23, 24, 25], synaptic receptor trafficking at neuronal synapses [26] and SVG virus assembly [12]. A significant recent advance is the analysis of massive numbers of overlapping SPTs. This statistical analysis can reveal the properties of molecular trajectories. However analyzing these data can potentially also allow quantification of membrane dynamics [27] and may give insight into organelle organization.

Here we have developed a method based on hybrid algorithms, and automated analysis pipeline for SPTs (Fig. 1C), based on the Langevin [28] equation of motion to provide statistical analysis of these data. This method estimates biophysical characteristics and is capable of reconstructing nanodomain sizes and boundaries using the classical physical model of a potential well [29, 28], well known since Kramer’s work in 1940. This method allowed us to characterize calcium channel organisation in the membranes of hippocampal neurons. We also present an algorithm that reconstructs a network from SPTs based on the clustering of low-velocity trajectory fragments, and use it to reconstruct ER network topology as well as estimate the time scale of lysosome trafficking and ER network interactions. Finally, these methods allowed us to extract the motion of trajectories relative to the reconstructed network, thereby revealing the redistribution of trajectories inside the ER of normal and atlastin-null mutant cells [30]. The diversity of datasets used here demonstrate the broad applicability of these methods to cell biological processes. The methods are available to users as elementary ImageJ plugins.

## 2 Results

### 2.1 An algorithm that reconstructs nanodomains in high density regions

In order to extract sub-cellular regions where a high density of molecular interactions are occurring, we develop a novel computational approach and automated pipeline Fig. 1C, based on stochastic equations, multiscale analysis, optimal estimators and maximum likelihood statistics. In contrast with methods to extract density of points [31, 32] or to compute the Maximum Likelihood Estimator (MLE) [33, 34], the present approach combines density of points with local dynamics associated with the elementary displacement Δ*X* = ***X***(*t* + Δ*t*) — ***X***(*t*), where ***X***(*t*) is the position of the particle at time *t* [13, 14] to extract the field of forces. We could thereby assess the nature, organization and stability of a large amount of HDRs, by collecting statistics that can reveal hidden cellular organization. HDRs could previously only be characterized by manual curation based on extracting parameters of potential well, a concept in classical physics [35, 28] that describes the stability of dynamic systems, such as the motion of a bead attached to a spring. In contrast, using combined optimization methods, the present method allows users to automatically extract local diffusion coefficients, energetics, local field potential, and most importantly, local boundaries.

The method relies on an interesting observation that nanodomains of high density tend to have an elliptic shape. Thus the first step to detect them is to recover their center and boundaries (see method and eq. 12). Non-automated classification algorithms have used the density of points [33] or displacements (Δ*X*) separately. But these procedures often lead to parameter estimations that are not completely satisfactory due to a shallow minimum of the associated error function (see Method). This shallow minimum leads to large variability and possible mistakes in the estimation of most of the nanodomain parameters such as the boundary and the energy of nanodomains.

To overcome these difficulties, we developed a hybrid algorithm (Fig. 1C), described in the Methods section, which combines two independent procedures starting with a principal component analysis (PCA) to recover the elliptic boundary and followed by a MLE to compute the effective diffusion coefficient and drift properties.

To test this hybrid algorithm, we first constructed a ground truth dataset. This dataset consists of trajectories (Fig. S1 Aa-b) generated by the stochastic equation 2, where the force defining the nanodomain is defined by equation 12 and characterized by converging arrows (Fig. S1 Ac). We use a time discretization Δ*t* = 20 and 50 ms and generated 10^3^ trajectories of size 20 points. The new algorithm comprises three steps:

1. Automatic determination of bins with the highest-density of points (Fig. S1 Ba).
2. For each such bin, we iterate over square regions of increasing width *w_k_* around the bin center. For each iteration *k*, we apply a PCA to estimate the semi-axis *a_k_, b_k_*, the center *c_k_* and a score 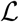 (eq. 24) based on the points falling in the square of size *w_k_* (Fig. S1 Bb-c). We iterate until we reach a threshold size defined by the user, as the maximum size of a well (Fig. S1Bc).
3. In the termination step, we select the optimal value of the iteration providing the optimal parameter (see Method paragraph 6.3).

The reconstruction is illustrated in Fig. S1C for three wells (see also Table S1 and S2 where we used two time steps Δ*t* = 20 and 50 ms to test the reconstruction of circular and elliptic boundaries). Finally, we showed that this hybrid algorithm performs much better than two previous algorithms in estimating all parameters, especially those at low energy at 4kT (Table S1). Other algorithms are based on the local density of points [33], or on computing the drift term [14] only.

At this stage, we have thus validated the hybrid algorithm by using a groundtruth data sets to identify and automatically reconstruct nanodomains. Our novel hybrid algorithm out-performed existing models for all parameter estimations and in terms of defining the exact shape of the nanodomain.

### 2.2 Analysis of SPTs for the endogeneous voltage-gated calcium channels reveals their organization and weaker stability in nanodomains

We applied the hybrid algorithm to study the dynamics of calcium voltage channels (CaV2.1) tracking on the surface of neuronal cells for two overexpressed splice variants CaV2.1_Δ47_ and CaV2.1_+47_, previously shown to shape synaptic short-term plasticity [14]. We also examined endogenously tagged CaV2.1 channels. Using a large number of redundant SPTs (see Methods), we were able to automatically detect the nanodomains defined as HDRs. We plotted the trajectories, density, diffusion maps and the drift field associated with these HDRs (Fig. 2A-C). The algorithm allowed us to identify the geometrical characteristics of nanodomains approximated as ellipses with semi-axis lengths *a* = 143 ± 51 nm and *b* = 104 ± 33 nm for Cav2.1_Δ4_7. These parameters are similar for Cav2.1_+47_ (Fig. 2D), while it was smaller for endogenous Cav2.1 with *a* = 100±47 nm and b = 73±31 nm. The diffusion coefficient was *D* = 0.091 ± 0.052 *μm*^2^/*s* for the Δ47 variant, *D* = 0.087 ± 0.045 *μm*^2^/*s* for the +47 variant and smaller for endogenous CaV2.1 with *D* = 0.069 ± 0.051 *μm*^2^/*s*.

**Fig. 2.**
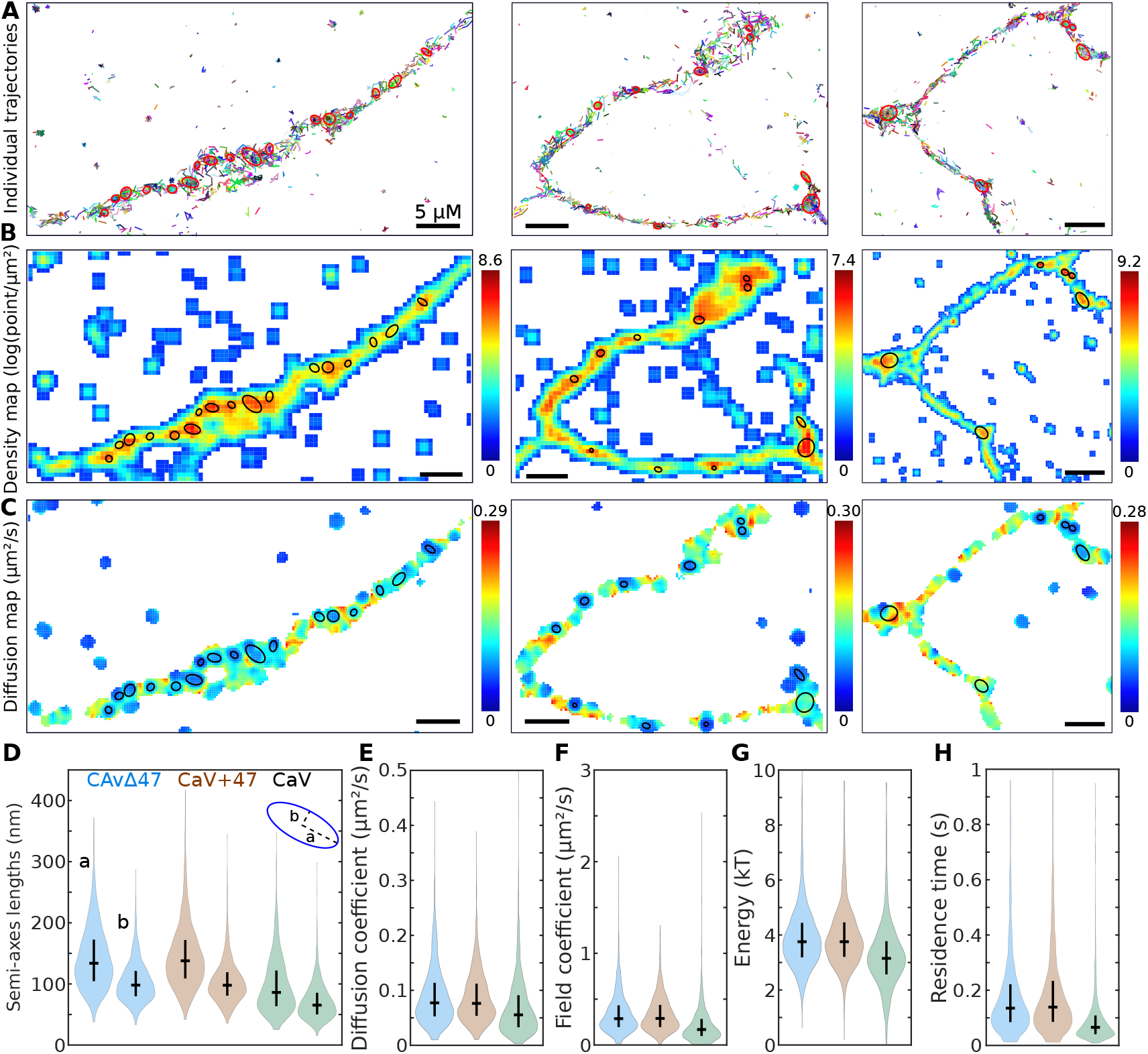
Large statistics of CaV nanodomains located on dendrite automatically detected. **Super-resolution SPTs automated analysis reveals CaV nanodomain organization. A.** Examples of trajectories displaying high-density regions (red ellipses). **B.** Associated density map, presenting the local point density (in log(points/*μm*^2^)) and computed over a grid with bin sizes of 50 nm and locally averaged over a 3×3 Gaussian kernel. **C.** Associated diffusion map, displaying the local diffusion coefficient (*μm*^2^/*s*) computed over a grid with bin sizes of 80 nm with at least 15 displacements and estimated with the cosine-round method (Material and Method) with a radius of 100 nm. **D-H.** Population characteristics for three experimental conditions: tracking CaV2.1-47, CaV2.1+47 or CaV2.1. Nanodomain represented as potential wells are recovered using the hybrid algorithm. D: Distribution of semi-axes lengths of elliptic well boundaries, E: Diffusion coefficients inside wells, F: Field coefficients *A*, G: Energies of the well (kT), H: Residence time distribution of a trajectory inside a well (H). Mean and variance values are shown in Table S3.

Interestingly although the associated energy was ~ 3.9 kT for the two variants and 3.3 kT for endogenous CaV 2.1, we found that the associated residence times of a receptor in a nanodomain was around 174 – 181 ms and 94 ms respectively (Table S3). We also tested two methods to estimate the diffusion coefficient, either using a static grid [33] or a sliding disk convolved with a cosine function (Fig. S1) and used this approach in subsequent analyses. To conclude, these estimations of residence time were larger (almost double) compared to the ones we reported in [14] and thus our new approach quantifies channel dynamics in the neuronal membrane more precisely. The approach also captures differences in variant forms of channels, revealing a significant reduction in interactions for the endogenous fraction.

### 2.3 Time-lapse analysis reveals the stability of CaV nanodomains over time

To investigate the stability of nanodomains across time, we used a timelapse analysis (Fig. S3) with sliding windows of 20 s and no overlap. This analysis allows determination of the lifetime of a nanodomain, which is given by the number of successive windows where it is detected (Fig. S4 A). For example, the trajectories obtained during a 250 s experiment are split into 13 20 s windows (0 – 20 s, 20 – 40 s, …, 240 – 260 s). We searched for the presence of potential wells in each window (Fig S4A-B). To follow a well across successive time windows, we consider that two wells identified at time *t_k_* and *t*_*k*+1_ are the same if the distance between their centers is less than 250 nm. The ensemble of consecutive times (*t_q_*,…*t_r_*) where a well is first detected at time *t_q_* and disappears at time *t*_*r*+1_ is used to define the stability duration *τ* = *t_r_* – *t_q_*. This analysis allows us to follow the size of the small and large axis of the wells and the associated energy over time (Fig. S4C-D). Finally, we found that 55% of the wells were present for more than 20 s (one time window) and that their average duration is ~ 56 s for both Δ47 and +47 variants (Fig. S4E and Table S4). However, nanodomain stability is reduced to *T* = 46 s for endogenous CaV2.1. All of these durations are longer than the ~ 30 s that we previously reported [14], indicating that the present algorithm advances our ability to capture the dynamics of these high density enigmatic subcellular domains.

### 2.4 The hybrid algorithm reveals that ER node are nanodomains defined by an attracting field of force

To further explore the range of applicability of the present hybrid algorithm, we analysed SPTs recorded from the ER of HEK293T, COS-7 WT as well as atlastin knockout COS-7 dATL cells. In such mutant, the morphology of peripheral ER tubules is altered but it is unclear how the ER flow is affected. Since nodes have been previously characterized as HDRs [17], we asked here whether these nanodomains could further be defined as potential wells. Applying the hybrid algorithm reveals several potential wells (Fig. 3A-B red ellipses) precisely located at nodes forming high density regions. We further estimated the diffusion and (Fig. 3C) drift maps, observing converging arrows patterns in these regions (Fig. 3D and Fig. S4-A), a classical feature of potential wells [22]. Interestingly, these HDRs were characterized by ellipses with a large semi-axis *a* = 219 ± 71 nm and *b* = 155 ± 56 nm for HEK293T (see also Fig. 3E and table S6 for COS-7 WT and COS-7 dATL). The hybrid algorithm further reveals an average energy of *E* = 3.3 ± 0.9 kT and a mean residence time of *τ* = 101 ± 64 ms. Interestingly, although the elliptic parameters are not much different in the case of COS-7 WT and COS-7 dATL, a difference can be observed in the dynamical parameters characterizing the transport of the material across the ER network. To conclude, the present hybrid algorithm reveals that ER nodes concentrate trafficking of luminal molecules by a spring-force type mechanism, the origin of which should be further explored.

**Fig. 3.**
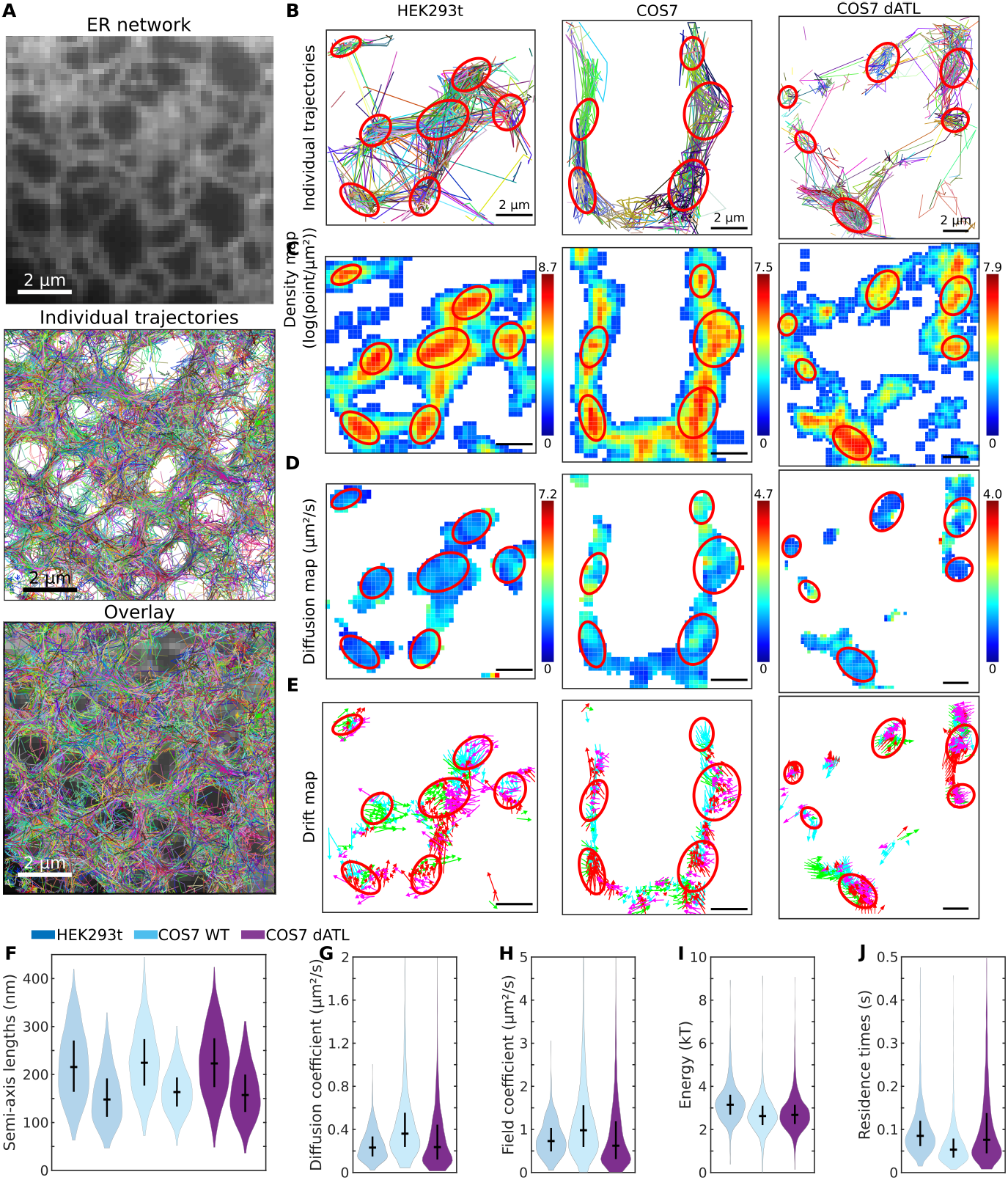
HDR present in organelle such as the ER network. **ER nodes revealed by super-resolution SPTs are characterized as potential wells. A.** Super-resolution ER network. Single trajectories recorded in the lumen and coded by a given color (second) and overlay (Third) **B.** Single trajectories coded by a given color and recorded inside the ER for three different cells (HEK293t, COS7, COS7-dATL). HDRs are associated with ER node (red ellipse). **B-C-D.** Density, Diffusion and Drift maps for the regions shown in **A.**. Arrows in the drift maps are colored according to the direction: West (purple), East (green), South (blue) and North (red). **E-J.** Statistics of the wells reconstructed from the hybrid algorithm (Material and Method): semi-axis (longest *a* and shortest *b*), diffusion coefficients *D*, Field coefficient *A*, Energy of the well *E* and the estimated residence time.

## 3 Organelle network reconstruction from a large number of SPTs

We next wanted to check whether our approach can be used to define the structure of the ER and be generalized to other organelles. For these goals, we developed a novel method and algorithm to reconstruct the network from SPTs.

### 3.1 Graph Reconstruction Algorithm (GRA) to unravel the ER network

Although SPTs can be used to explore the ER network architecture [17], we still lacked a method for automated reconstruction of the ER. We therefore developed the algorithm to see if the dynamic architecture of this complex organelle can be recovered from SPTs.

The starting point is to colour trajectory displacements depending on their instantaneous velocity. This reveals a dynamical segregation of the ER into nodes and tubules (Fig. 4A-C). Based on this segregation, we developed the Graph Reconstruction Algorithm (GRA) to recover the ER structure from SPTs. The GRA consists of three steps: 1-identifying the HDRs formed by low-velocity (blue regions) trajectory fragments forming the nodes of the graph (Fig. 4D-G), 2-determining the high-velocity trajectory fragments connecting the previously detected nodes (Fig. 4H-K) and 3-constructing the associated graph (Fig. 4L), as described in Materials and Methods. The first step of the method relies on the density-based clustering algorithm (dbscan) [36], which requires that the distribution of trajectories be quite heterogeneous so that small regions of high density are well-separated. In an additional step, a direction can be attributed to the links (i.e. tubules) based on the percentage of trajectories going in the same direction between two nodes: when the percentage is around 50%, the node is bidirectional, however if the percentage is much lower, the node is defined as unidirectional. This procedure allows us reconstruct a two-dimensional graph for the organelle network that can be used to study further statistical properties.

**Fig. 4.**
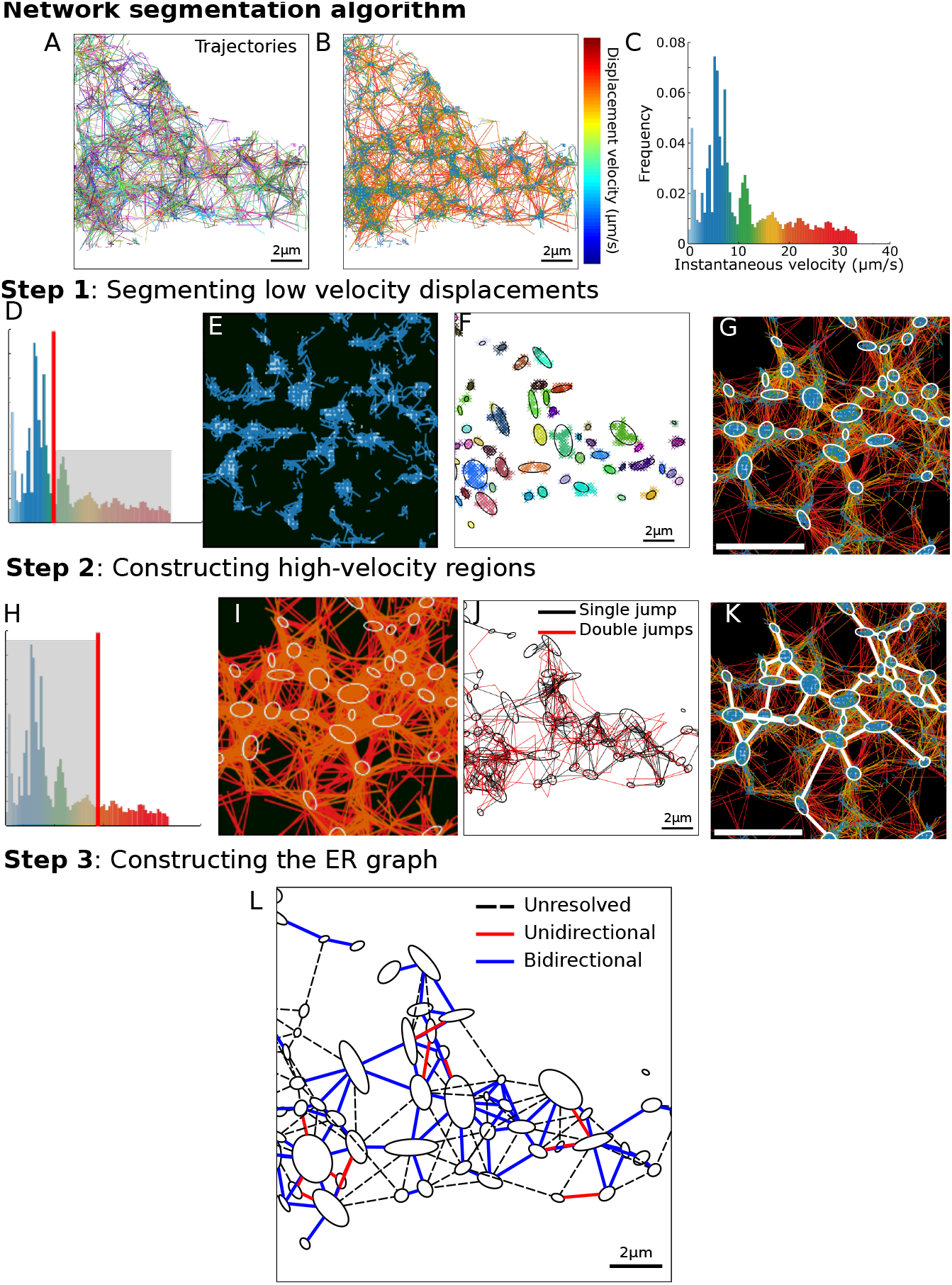
Algorithm to automatically reconstruct a network. **Automatically network segmentation revealed by super-resolution SPTs. A-C.** SPTs (A), density map (B) and color coded distribution according to amplitude of instantaneous velocity (C). Step 1: detecting regions of low velocities. **D.** dbscan procedure to cluster (E) and merge them into ellipses (F) and embedding into the network (G). Step 2: characterization of high velocity regions from the histogram (H), segmented between ellipses (step 1) showing one or two fast jumps (J) and finally reconstructed in the ensemble of trajectories (K). Step 3: abstract graph reconstruction of an ER showing nodes and tubules, that could be bi- or unidirectional. Step 4: Synchronizing trajectories reveal a switching dynamics occurring inside tubules.

### 3.2 Graph Reconstruction Algorithm of the ER network of COS-7 dATL cells reveals aberrant organization and trafficking

Next we examined whether disruptions to organelle structure and dynamics can be captured using our algorithm. Atlastin-1 is a GTPase that mediates homotypic membrane fusion in the ER. We analyzed SPTs recorded from an ER luminal probe in atlastin knockout (dATL) COS-7 cells. We present colour-coded trajectories that follow their instantaneous velocity (Fig. 5A-B), the density and diffusion maps (Fig. 5C-D), as well as the histogram of the apparent diffusion coefficients (Fig. 5E), revealing a mean of *D_app_* = 1.56 ± 0.83 *μm*^2^/*s*.

**Fig. 5.**
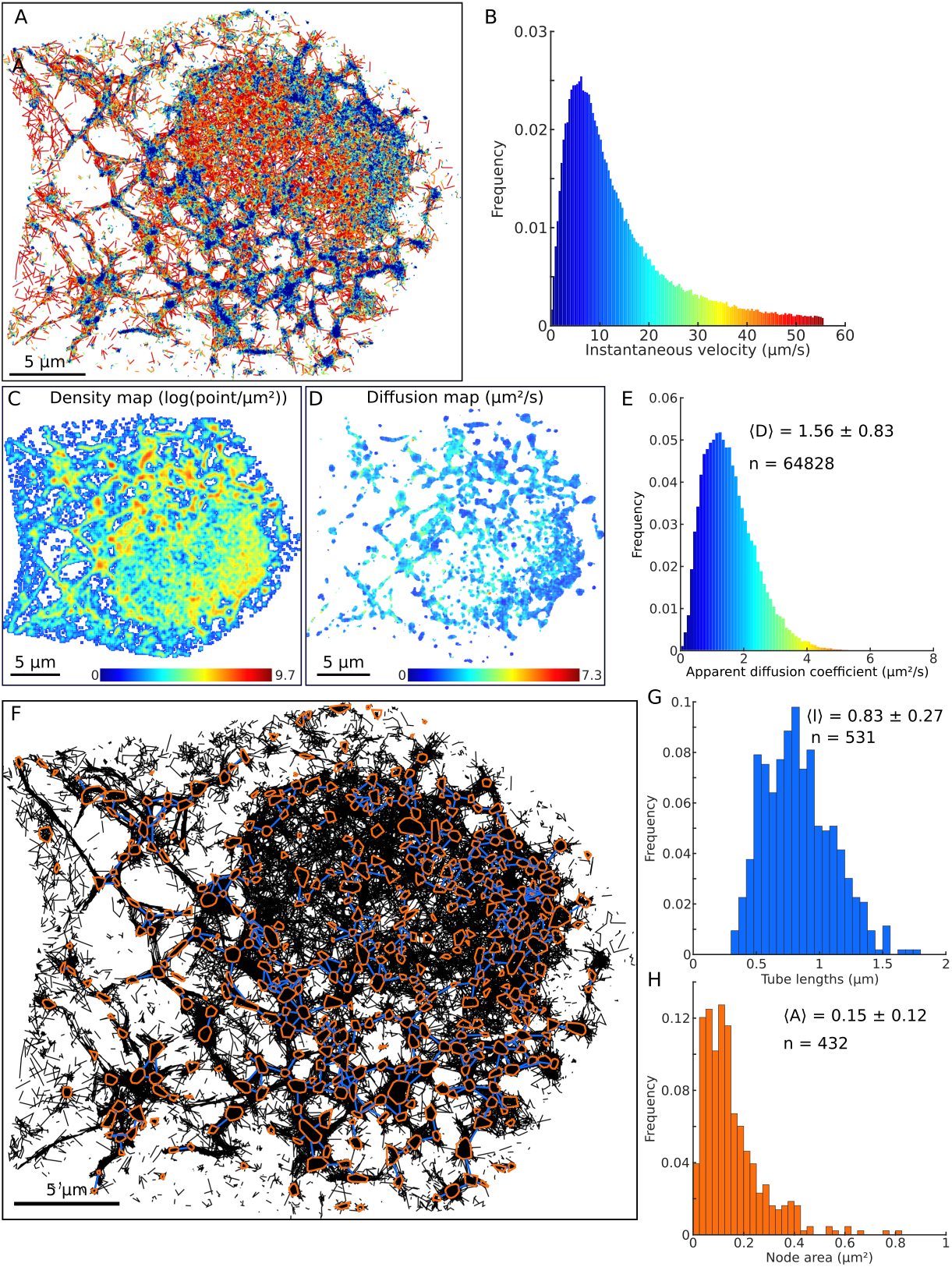
ER network in dATL cells. ER network reconstructed in COS-7 dATL cells. **A** Individual trajectories color-coded according to their instantaneous velocity shown in **B**. **C** Density and **D** drift maps. **E** Distribution of diffusion coefficients obtained from the individual bins of the diffusion map presented in D. **F** Reconstructed network, showing the nodes in red and links in blue overlaid on top of the individual trajectories (black). **G** Distribution of distances (i.e. tubule lengths), between connected nodes. **H** Distribution of the areas covered by nodes.

We applied the GRA and obtained a graph reconstruction of the ER (Fig. 5F) where the HDRs (red) are connected by blue segments. This approach allows quantification of the distribution of distances between nodes with a mean *d_nodes_* = 0.83 ± 0.27 *μm* (Fig. 5G) and the node projected area *S_nodes_* = 0.15 ± 0.12 *μm*^2^ (Fig. 5H). Note that the RGA could miss some non-explored ER regions or regions that are sub-sampled. To conclude, the GRA allows automated reconstruction of organelle networks based on SPT exploration once there is enough heterogeneity in the distribution of datapoints.

### 3.3 Graph Reconstruction Algorithm and SPTs segmentation reveal the duration of ER-lysosome interactions

To demonstrate the broad applicability of the algorithms, we analyzed an ensemble of trajectories from lysosomes (Fig. 9.6A). These trajectories are characterized by a distribution of instantaneous velocities in the range [0 – 3.5] *μm/s* (Fig. 9.6B). However low and high velocities are not segregated, but are instead found in similar regions (Fig. 9.6A left and right). The distribution *f*(*v*) of velocities (Fig. 9.6B), can be fitted by a sum of two exponentials

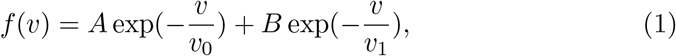

where a best fit approximation gives *v*_0_ = 0.06 *μm/s* (95% confidence interval [0.057, 0.072]) and *v*_1_ = 0.6 *μm/s* (95% confidence interval [0.383,1.322]) (coefficient of determination *R*^2^ = 0.95), with *A* = 0.20 and *B* = 0.013. This fit suggests that the distribution of lysosomes is largely driven by low velocity components. The rare appearance of high velocity components suggests a possible switch between slow and fast motions. Finally, note that 86% of displacements are associated with a velocity of less than 0.5 *μm/s* and 12.9% are in the range of [0.5-1.5] *μm/s.*

**Fig. 6.**
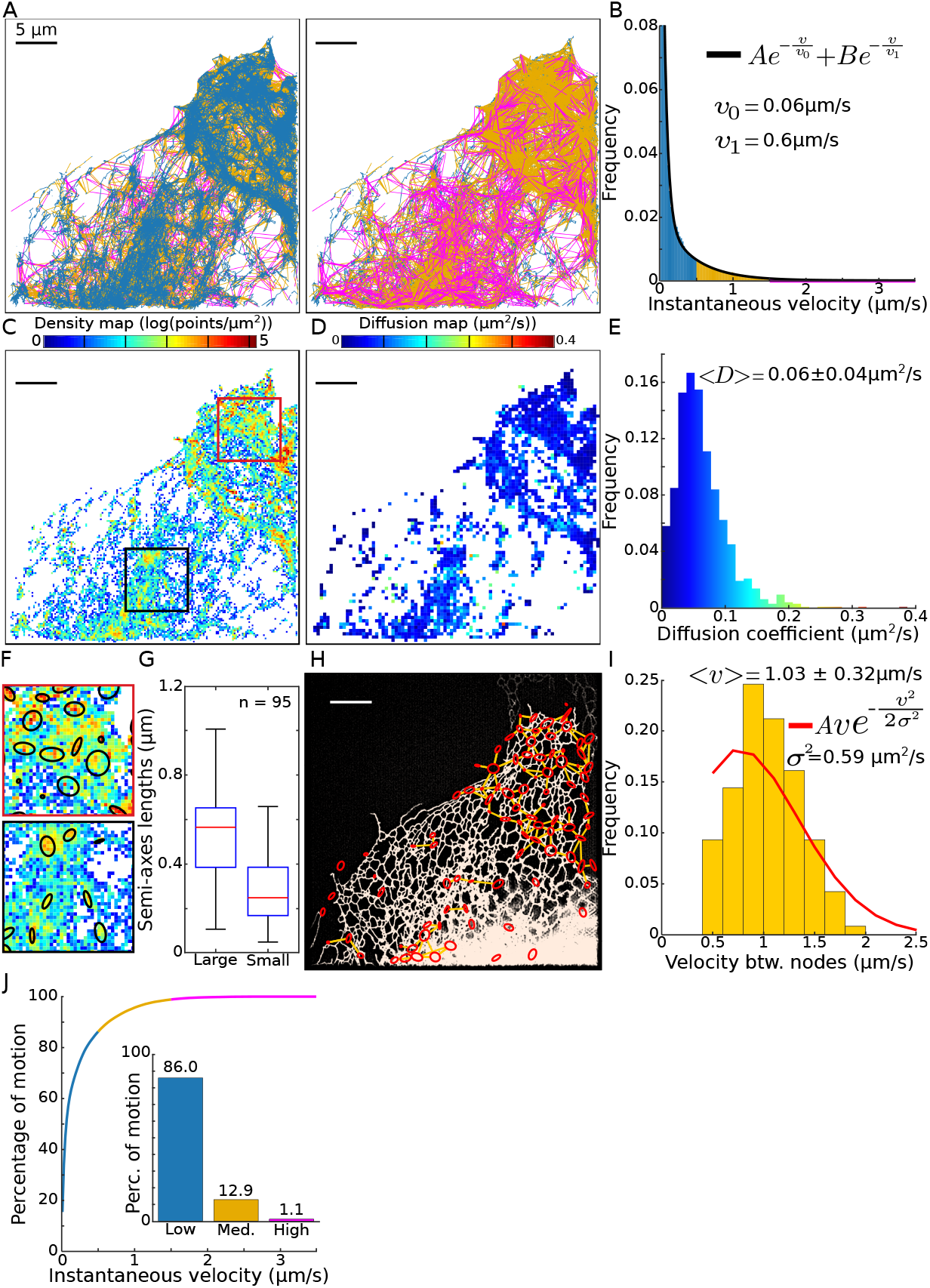
Reconstructing ER network from lysosome trajectories. Lysosome trajectories analysis. **A**. Lysosome trajectories color-coded according to individual displacement amplitudes (panel B). **B**. Instantaneous velocities, color-coded with respect to the value and fitted by a double exponential distribution. **C**. Density map of points. **D**. Diffusion map presenting the local diffusion coefficients. **E**. Histogram of the diffusion coefficients obtained in the square bins form panel *D*. **F**. Magnification of the density map of two regions of interest, showing high-density regions, approximated by ellipses. **G**. Length of semi-axes of high-density regions approximated as ellipses. **H**. Reconstruction of a lysosome graph, where nodes correspond to high-density regions. A link (in yellow) is added when at least one trajectory starts in one node and enter to the other one (in one or two frames).**I**. Average instantaneous velocities between pairs of connected nodes found in panel F. **J**. Percentage of displacements with a specific instantaneous velocity. Inset, percentage of displacements for the velocity regimes defined in panels A-B.

To further study how lysosomes move in the cytoplasm, we computed relevant density and diffusion maps (Fig. 9.6C-D) and found that the motion had a diffusion component (Fig. 9.6E), with an average apparent diffusion coefficient of *D_app_* = 0.062 ± 0.040 *μm*^2^/*s*. Interestingly, regions of low diffusion coefficients colocalized with regions of high density in the density map [37, 27] (Fig. 9.6A-C).

We then isolated regions of high density using a method based on the density of points (Methods subsection 6.8.2), revealing an ensemble of *n* = 95 HDR sub-domains, approximated by ellipses (magenta in Fig. 9.6F) of semi-axis lengths *a* = 516± 196 nm (large) and *b* = 278± 143 nm (small) (Fig. 9.6G). By considering the displacements connecting different regions, we reconstructed (Method subsection 6.8.2) a network explored by the lysosomes, Fig 9.6H, where high density regions (red circles) are connected by direct lines (yellow). Interestingly, the histogram of average velocities between these regions is not symmetric (Fig. 9.6I) with a mean velocity *v* = 1.03±0.32 *μ*m/s, which clearly deviates from diffusion, as computed from the Rayleigh distribution. This deviation suggests that the transitions between these regions are driven by an active motion. Moreover, the overlay between ER (white) and the lysosome reconstructed network (Fig. 9.6G) suggests that the lysosome trajectories follow the topology of the ER network [7]. To conclude, our analysis reveals that lysosomes travel along a network that strongly colocalizes with the ER. However, high and low velocities occur in similar regions. Since lysosomes move along microtubules, this present statistical analysis suggests that the ER-microtubule network shapes lysosome trafficking.

## 4 Trajectory re-synchronization approach reveals single local molecular dynamic exploration inside an ensemble

In the previous result sections, we reconstructed the networks hidden inside SPTs data. We shall now introduce a last step in our method which resynchronizes trajectories that fall inside the same subcellular are, but were acquired at random times. This approach allows us to study the dynamics of trajectories with respect to the ensemble of trajectories that visit the same spatial region. The approach also enables determination of the local spatio-temporal properties that trajectories explore at the single unit level.

### 4.1 Trajectory segmentation reveals ER-lysosomes interaction time scale

To study the possible interactions between lysosomes and the ER (Fig. 9.7A) reconstructed network in Fig. 9.6, we focused on the confined portion found along individual trajectories (see method section 6.8.3 and Fig. 9.7A). We hypothesise that the lysosome motion can switch between a directed and confined motion (Fig. 9.7B). We first show that the lysosome can indeed switch between directed and confined motion (Fig. 9.7C).

**Fig. 7.**
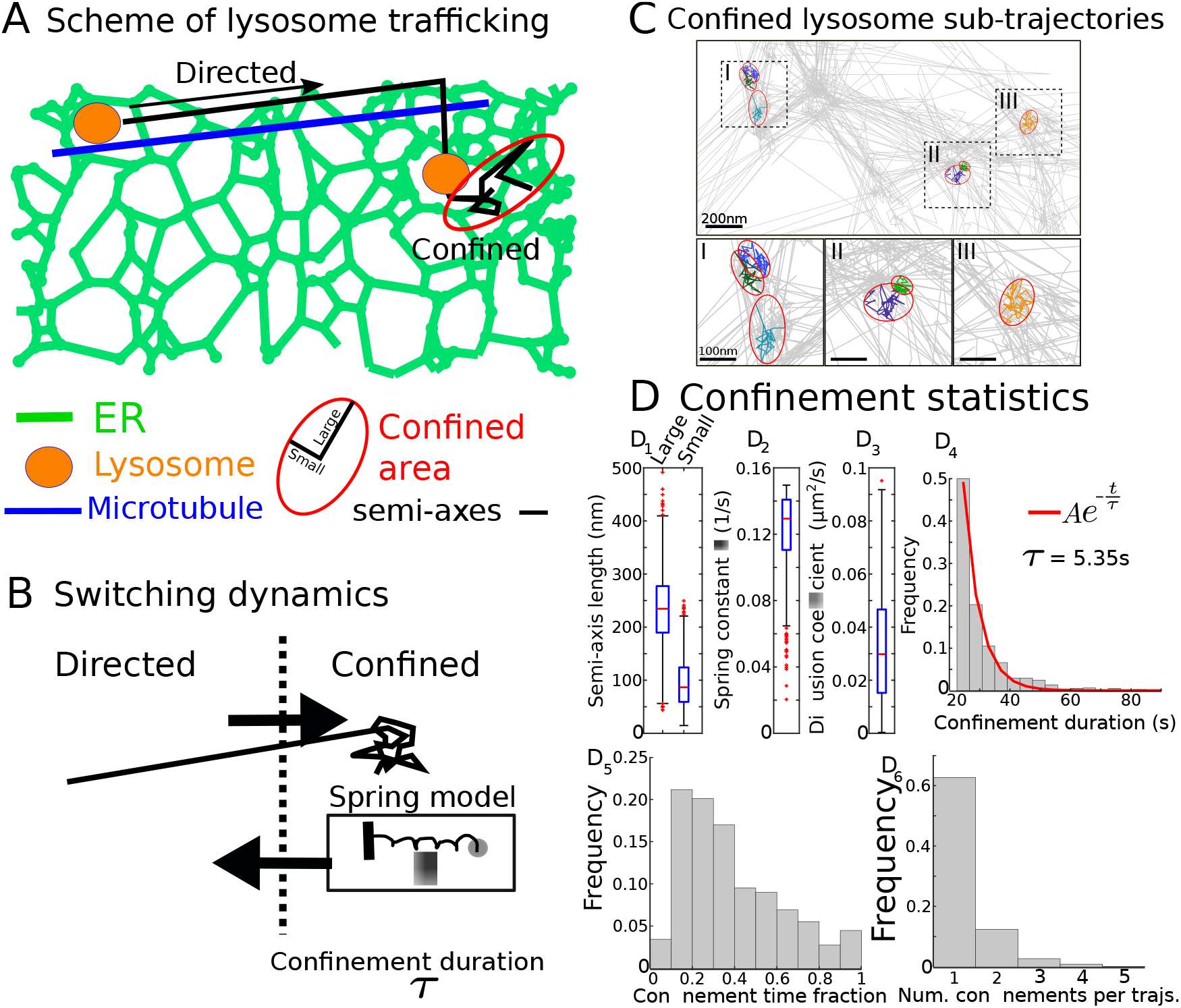
ER network-Lysosome interaction revealed by confined trajectories. Lysosome confinement time **A**. Schematic representation of a lysosome switching between a directed motion along microtubule and confined motion at ER nodes. **B**. Switching dynamics representation: the confined state is characterized by a spring constant λ a diffusion coefficient *D* and a confined time constant *τ*. **C**. Regions of confinement for individual lysosome trajectories: three examples (insets). **D**i. Semi-axes (small and large) of the ellipse fitted to the confinement regions. **D**_2_. Spring constants of an Ornstein-Ulenbeck process–confined diffusion–. **D**_3_. Diffusion coefficients estimated inside a confinement region. **D**_4_. Residence times inside a confinement region. **D**_5_. Fraction of time each trajectory spends confined (relative to the trajectory length). **D**_6_. Number of confinement events along individual trajectories.

To recover the size of the confinement areas, we fitted ellipses over these regions and obtained average semi-axes lengths (Fig. 9.7B) of *a* = 232 ± 77 nm (large) and *b* = 94±47 nm (small). Furthermore, this approach allowed us to estimate the confinement strength λ by considering that the confined motion could be generated by a spring force, modeled by an Ornstein-Uhlenbeck process [28]. We found that the spring force is λ = 0.123±0.025 *s*^-1^ (Fig. 9.7C), associated with an average local diffusion coefficient of *D* = 0.032 ± 0.002 *μm*^2^/*s* (Fig. 9.7D) for a total of *n* = 818 confinement regions. Finally, the distribution of times in confined regions could be well-approximated by a single exponential with a time constant *τ* = 5.35 s (Fig. 9.7E). The average residence time of lysosomes in these regions is 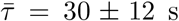 which can be interpreted as a time where lysosomes could interact with the ER. To conclude, the present algorithm reveals that as lysosomes travel along a network that strongly colocalizes with the ER, the velocity can switch from large to small displacements and the trajectories can become restricted into regions of size ~ 200nm, on a timescale of 5 s. This could correspond to a change in directionality of movement or a direct interaction with the ER. Our approach can therefore extract changes in lysosome dynamics that may reflect functional interactions from complex data.

### 4.2 Trajectory re-synchronization approach shows how a single trajectory explores single nodes

After an ER network is reconstructed from SPTs by the GRA (Fig. 9.8A), the node-tubule topology emerges. Thus it becomes possible to study how trajectories locally explore the network by synchronizing them upon exit from a chosen node (Fig.9.8B). Interestingly, we found that the mean instantaneous velocity at exit is *v_e_* = 30.2 ± 10.2 *μm/s* and keeps decreasing during the next 200 to 300 ms (Fig. 9.8C). Escape occurs in equal directions (Fig. 9.8D), as shown in four examples where we followed their dispersal. To characterize this dispersion, we plotted the dispersion index (section S5), revealing two phases (Fig. 9.8B), one below 100ms, showing a rapid dispersion, followed by a second phase with less expansion. These two phases can be interpreted as: for the first one, trajectories escape from a well, then in the second phase, trajectories tend to be re-captured for a certain time in nodes, thus preventing a fast exploration of the network.

**Fig. 8.**
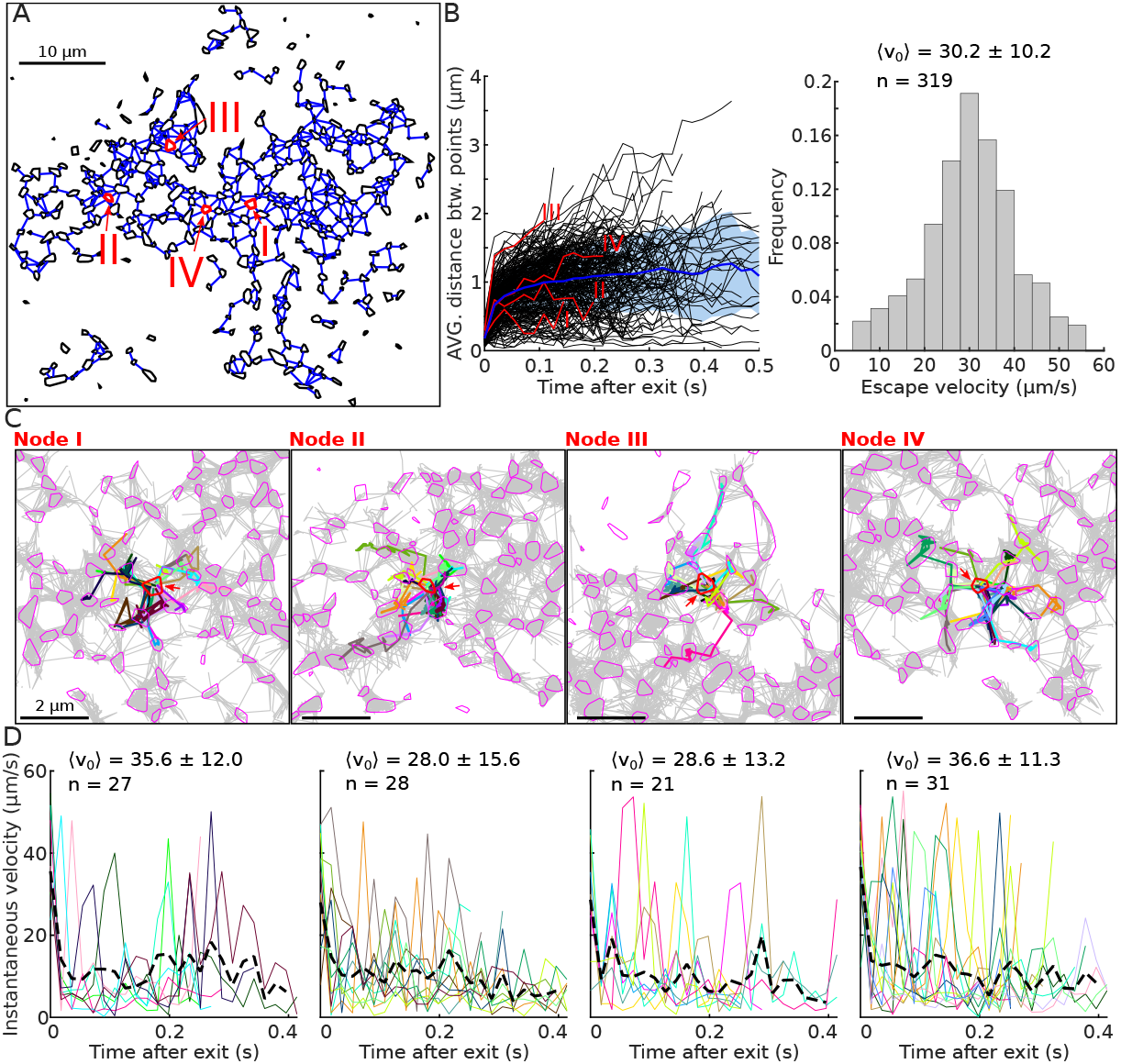
Local space exploration after trajectory re-synchronization. **Local network exploration revealed by spatial re-synchronization of superresolution SPTs. A.** ER reconstructed network connectivity using the GRA with nodes (black polygons) connected by segments (blue). Four nodes (I, II, III, IV) are selected (red arrows) inside a ER segmented network. **B.** Average distance between exit points vs time after escape. We highlight 4 examples (red), average (blue) and individual examples (black), fast (< 50*ms*) and a slow phase. **C.** Local exploration of neighboring nodes from trajectories (various colors) exiting from the chosen node.**D.** Distribution of velocities for trajectories exiting a node for the four examples: the mean velocity decreases from a high amplitude corresponding to the instant of exit.

### 4.3 Trajectory re-synchronization reveals novel local dynamics within dATL ER-tubules

We next analyzed SPTs recorded in the ER of COS-7 dATL cells which lack the ER membrane-shaping protein atlastin (double knockout of the ATL-2/3 genes [30]) and exhibit a disrupted peripheral ER morphology with elongated tubules [30]. These SPTs revealed changes in the local space exploration of trajectories (Fig. S7A). Under normal conditions, trajectories are mostly located in nodes [17], while here trajectories predominantly explore long tubules (fig. S7B) with lengths of 5.4 ± 2.44 *μm*, much longer than the ~1 *μm* found for the tubules of WT cells. Inside these tubules, we found that trajectories exhibit a ”stuttering” behavior around different positions that lasts for seconds. To characterize this behavior, we estimated several parameters such as the duration that a trajectory spent around a given position *τ_u_* = 79 ± 76 ms (Fig. S7D), the transition time between different positions *τ_t_* = 27 ± 15 ms (Fig. S7E), the length of a transition step Δ_*ll*_ = 0.53 ± 0.45 *μm*(Fig. S7F) and finally the standard deviation around the stable positions SD = 0.14 ± 0.07 *μm* (Fig. S7G).

To conclude, following the reconstruction of networks using our algorithm we were able to re-position and re-synchronize SPTs. Using analyses of the ER-lysosome, ER in normal COS-7 cells, and ER in COS-7 dATL cells, the algorithm revealed trajectories explored by the local geometrical space and the associated time scales.

## 5 Discussion and concluding remarks

We present here a general method and the associated algorithms that can automatically characterize nanodomains where trajectories accumulate. Our approach generate graph representations organelle networks from SPTs. Automatically finding nanodomains is useful to extract large statistics (size, energy of potential wells, mean residence time of particles) and compare their properties. Further, by quantifying the trajectories inside and outside HDRs, we could recover membrane organization, as well as determine the local redistribution of dynamic organelles and proteins. By reconstructing a graph of ER or lysosome networks from SPTs we can recover molecular flow at the nanoscale level. We found here that HDRs could be characterized as attractive nanoregion (potential wells), and this generic representation suggests a universal mechanism of molecular stabilization that probably requires further investigation. Interestingly, these structures can be transiently remodeled in time as revealed by the present time-lapse analysis.

### 5.1 Universality of high density nanoregions characterized as potential wells

High density nanoregions are now associated with potential wells for several receptors and channels such as CaV [14], AMPAR [13], Glycine receptors [38] or GAGs [39, 40]. Interestingly, nodes of the ER can also be characterized as potential wells, which may reflect retention of luminal flow or to allow protein maturation. This representation suggests a generic membrane organization to retain particles (receptors, channels, proteins,etc…) in a field of force with long-range interactions.

Interestingly, the geometry of these regions and their energy profile are independent of the experimental conditions, further confirming again their stability. Note that the physical nature of potential well remains unclear [41]. The present method could be applied to analyse molecular crowding and the dynamics of nanodomains, thus clarifying processes relevant for phase separation at synapses [42]. With the development of new labelling methods, improved fluorophores and the ability to tag endogenous populations of molecules via CRISPR/Cas, it will soon be possible to investigate phase separation at a population level and with SPT, to track endogenous dynamics, offering novel opportunities for the present approach.

### 5.2 Trafficking in networks

The present method allowed us to reconstruct the ER network. The reconstruction algorithm reveals that lysosome trajectories follow ER network. This reconstructed network is further segregated into nodes and links, but low and high velocities are now much more mixed compared to the reconstruction obtained from luminal proteins. It is possible that lysosomes follow the cytoskeleton network which is correlated with the ER [7].

In addition, the distribution of lysosome velocity follows a double exponential (displacement histogram in Fig. 9.6) with fast (~0.6 *μm/s*) and slow (~0.06 *μm/s*) components. However, a more detailed analysis revealed that these velocities can be further subdivided into: 1) confined motion (Fig. 9.7) characterized by a residence time of ~ 5 s. 2) deterministic motion between HDRs (Fig.9.7 F-G), characterized by a distribution of velocity with an average of 1.03 *μ*m/s. It would be interesting to better characterize the switch between confined and rectilinear motion. Regions of deterministic velocities and those where diffusion can be found are often not well separated, suggesting that lysosomes can use various modes of transport, independently of the subregions where there are located. We found however some regions characterized by a high density of trajectories, with a low velocity, suggesting that there are possible trapping mechanisms to retain lysosomes in specific subregions of the ER, possibly at exit sites [43]. This mode of motion is quite different from the internal motion inside the ER lumen or on its membrane: in the first case, the node-tubule topology is associated with a diffusion-drift dynamics, while in the second case, the motion on the ER membrane is likely diffusion-based [17]. To conclude, the present analysis reveals that the confined time is around 5s, suggesting that during this time, lysosomes may be trapped in interaction with the ER.

Future work should reveal interaction times between Lysosomes and the ER. By applying our algorithm to different cells and organelles, we have shown that information on the boundaries and dynamics of subcellular interactions can be revealed from large SPT datasets. The automated algorithms presented here can be applied by users to analyse hundreds of thousands of trajectories and to study nanodomains with almost no human intervention and are available as an elementary ImageJ plugin.

## 6 Material and Methods

The method section is organized as several subsections describing the physical model to interpret SPTs, the associated data estimators, the potential well description, several hybrid algorithms to analyse automatically the regions of high densities and finally a method to reconstruct a graph for organelle networks.

### 6.1 Diffusion model, velocity, vector fields and empirical estimators

In the Smoluchowski’s limit of the Langevin equation [28], the position ***X***(*t*) of a molecule is described by

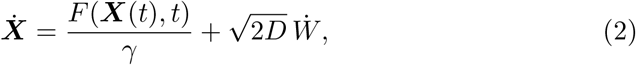

where *F*(***X***) is a field of force, *W* is a white noise, *γ* is the friction coefficient [44] and *D* is the diffusion coefficient. At a coarser spatio-temporal scale, the motion can be coarse-grained as a stochastic process [13, 37]

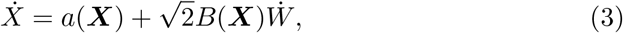

where *a*(***X***) is the drift field and *B*(**X**) the diffusion matrix. The effective diffusion tensor is given by 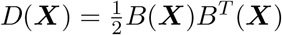 (.^*T*^ denotes the transposition) [45, 44]. The drift of the stochastic model from eq. 3 can be recovered from SPTs acquired at any infinitesimal time step Δ*t* by estimating the conditional moments of the trajectory displacements Δ***X*** = ***X*** (*t* + Δ*t*) – ***X*** (*t*) [44, 46, 47, 37, 22]

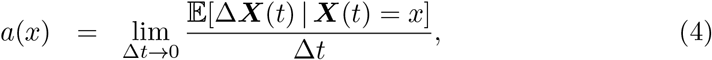

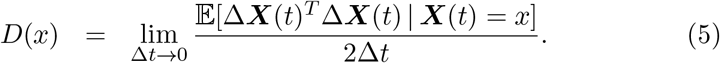

The notation 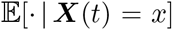 represents averaging over all trajectories that are passing at point *x* at time *t*. To estimate the local drift *a*(***X***) and diffusion coefficients *D*(***X***) at each point ***X*** of the membrane and at a fixed time resolution Δ*t*, we use a similar procedure to the one for the estimation of the density (section 6.4) based on a square grid.

The local estimators to recover the vector field and diffusion tensor [33] consist in grouping points of trajectories within a lattice of square bins *S*(*x_k_*, Δ*x*) centered at *x_k_* and of width Δ*x*. For an ensemble of *N* twodimensional trajectories 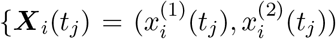, *i* = 1..*N*, *j* = 1..*M_i_*} with *M_i_* the number of points in trajectory ***X***_*i*_ and successive points are recorded with an acquisition time *t*_*j*+1_ – *t_j_* = Δ*t*. The discretization of eq. 4 for the drift *a*(*x_k_*) = (*a*^(1)^(*x_k_*), *a*^(2)^(*x_k_*)) in a bin centered at position *x_k_* is

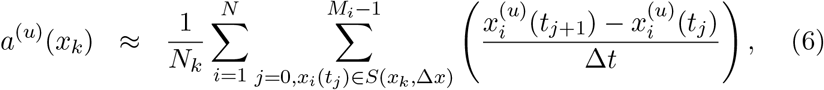

where *u* = 1..2 and *N_k_* is the number of points from any trajectory falling in the square *S*(*x_k_,r*). Similarly, the components of the effective diffusion tensor *D*(*x_k_*) are approximated by the empirical sums

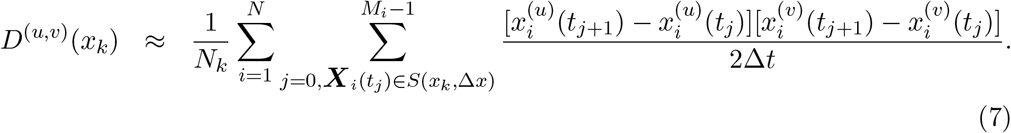

The centers of the bin and their size Δ*x* are free parameters that are optimized during the estimation procedure.

#### 6.1.1 Variant estimation of the diffusion coefficient and drift

To increase the accuracy of the diffusion and drift maps, we weighted the points in the moving windows with a cosine function (would also be possible to use wavelets). In that case, the new estimator for the drift field is now

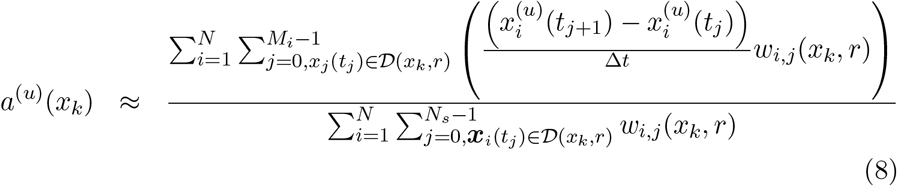

with *N_k_* the number of points of the trajectories falling in the disk 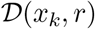 of radius *r* and centered at *x_k_*. The weight of a displacement starting at *X_i_*(*t_j_*) with respect to the disk 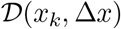 is given by

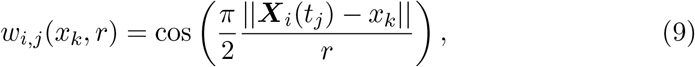

with ||.|| the Euclidean norm. In that case, we can choose a refined grid *S*(*x_k_*, (Δ*x*)′) with bin size (Δ*x*)′ = Δ*x/l_sc_*, where *l_sc_* is a scaling factor. The role of the cosine weights w is to decrease continuously the influence of the points falling near the boundary.

Similarly, the generalized formula for the effective diffusion tensor *D*(*x_k_*) are given by the weighted sums

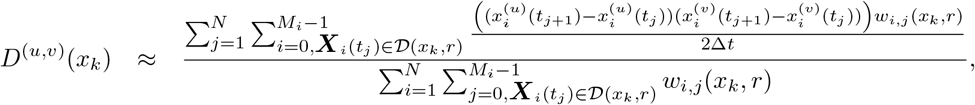

where the weights *w* are given by eq. 9.

#### 6.1.2 Local point density estimation

The local density of points *ρ* can be determined using a procedure similar to the drift or diffusion estimation by the image plane into a square bin *S*(*x_k_*, Δ*x*). We then compute for each square of *S* centered at *x_k_*

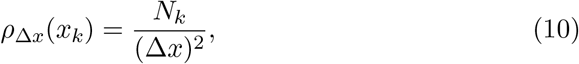

where *N_k_* is the number of trajectory points falling into the bin centered at *x_k_*. In practice, it usually helps to smooth this density estimation by applying a local average using a small *d* × *d* kernel with *d* ~ 1,3, 5.

### 6.2 Estimating potential well parameters

In this subsection, we present the estimators for the two main parameters of potential wells: the extent of their boundary and their associated energy [13, 38, 33]. We recall that the diffusion coefficient inside a well is considered to be constant and is computed from the second moment of the displacement for all points falling inside the boundary of the well (see eq. 4). The potential well model allows to estimate the residence time using the classical escape time formula [44, 13] for a circular well

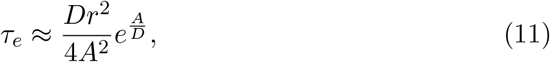

with *r* the radius of the well, *A* its attraction coefficient and *D* its diffusion coefficient. In the case of an elliptic well, we obtain an approximate circular boundary using the harmonic mean of the semi-axes 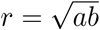, where *a* and *b* are the large and the small-axes lengths respectively. This approximation holds for *a* ≈ *b*.

#### 6.2.1 Parabolic potential well representation

To extract the energy of well, we consider the basin of attraction of a truncated elliptic parabola with the associated energy function

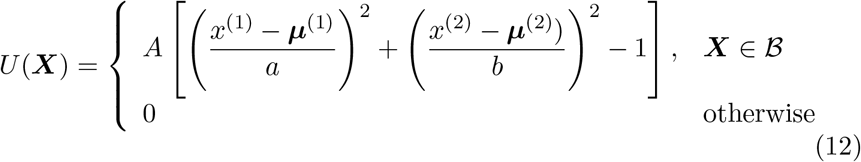

where ***X*** = [*x*^(1)^, *x*^(2)^], ***μ*** = [***μ***^(1)^, ***μ***^(2)^] is the center of the well, *a, b* are the elliptic semi-axes lengths and the elliptic boundary is defined by

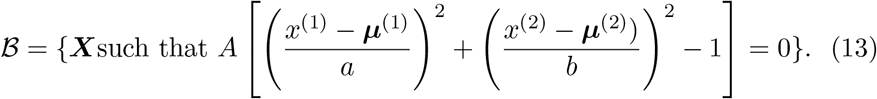

#### 6.2.2 Recovering the center *μ*

The center of the nanodomain 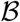 is estimated as the center of mass of the cloud of points falling inside the HDR. We use the empirical averaging formula

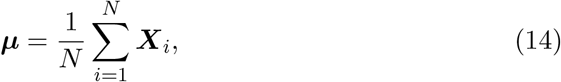

where *N* is the total number of points such that 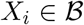.

#### 6.2.3 Covariance matrix Σ

We use the sample estimator of the Covariance matrix defined for a cloud of *N* two-dimensional points 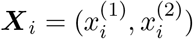 as

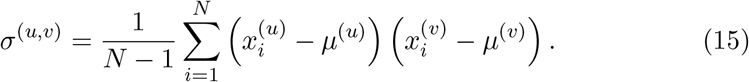

#### 6.2.4 Confidence ellipse estimation *ε* = (*μ, a, b, φ*)

We define the boundary of the well as the *X*% confidence ellipse of the associated Gaussian density distribution of center ***μ*** and covariance Σ. Using the singular value decomposition method, we decompose the covariance matrix Σ as

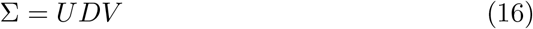

where *U, V* are unitary matrices and *D* is diagonal. The values in *D* represent the variance in each dimension along the principal components. The values of *D* follow a Chi-Square distribution with *n* = 2 degrees of freedom. Therefore the semi-axes lengths *a, b* can be obtained at the *x*% from *D* as

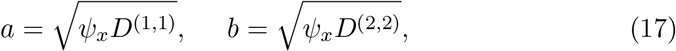

with *ψ_x_* is given by *P*(*v* < *ψ_x_*) = *x* for a Chi-Square distribution with two degrees of freedom (for example *ψ*_99_ = 9.210, *ψ*_90_ = 5.991 and *ψ*_90_ = 4.605). Finally, the orientation φ of the ellipse is defined by the angle

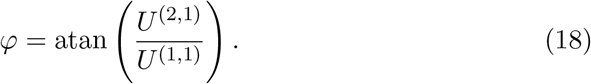

#### 6.2.5 Maximum likelihood estimators (MLE) based on an Ornstein-Uhlenbeck model

Using the potential well representation from eq. 2 in the stochastic model presented in eq.12 leads to a truncated Ornstein-Uhlenbeck process, centered at *μ*, with an attraction coefficient λ and diffusion coefficient 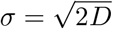. The probability density function *p*(***X*** (*t*_*j*+1_)|***X*** (*t_j_*)) for *j* = 1..(*M*–1) of observing two successive positions of the same trajectory ***X*** (*t_j_*) and ***X*** (*t*_*j*+1_), separated by a time step *t*_*j*+1_ – *t_j_* = Δ*t* is given by

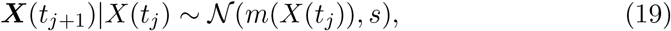

with

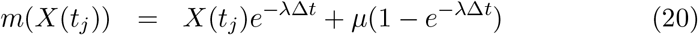

and

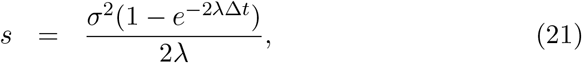

which we rewrite as

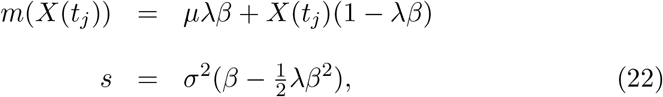

and

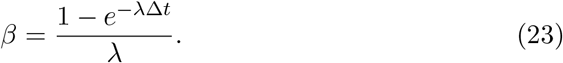

The log-likelihood function of observing the successive pairs (***X***_*i*_(*t_j_*), ***X***_*i*_(*t*_*j*+1_)), *i* = 1… *N*, possibly from various trajectories, is given by

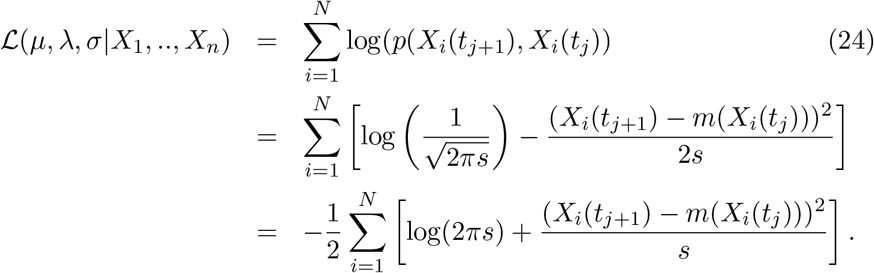

The corresponding maximum likelihood estimators for λ and 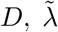 and 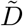 are obtained by solving the system of equations

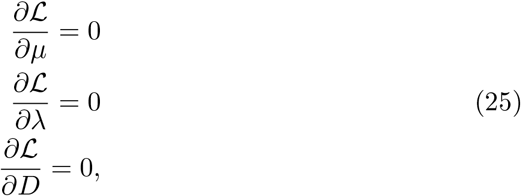

from which we obtain the empirical estimators for the drift coefficient

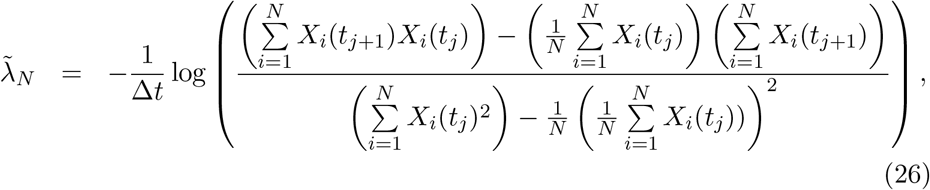

and the diffusion coefficient:

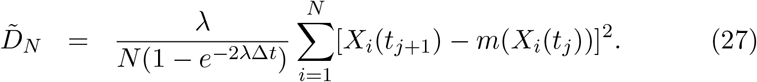

Note that the parameter λ in eq. 27 can be computed from the estimator 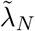.

### 6.3 Hybrid density-drift algorithm

In this sub-section, we present two variants of an algorithm to detect the main characteristics of a potential well from some observed trajectories: the center *μ*, the semi-axes lengths *a* ≥ *b*, the orientation *φ,* the field strength *A*, the diffusion coefficient *D* and the energy *E*.

#### 6.3.1 Fixed Spatial Scale Hybrid Density-Drift (FSHDD) Algorithm

##### Initiation

Search for high-density regions: the image is partitioned by a grid *G*_Δ*x*_ with square bins of size Δ*x*. from which we compute the density map *ρ*_Δ*x*_(*x*) (eq. (10)). We then select the bins from *ρ*_Δ*x*_(*x*) with the highest *d*% density as possible regions containing a potential well.

##### Iterations

For each region obtained in the initiation step, we apply an iterative procedure that is going to consider increasingly larger square neighborhoods around this region. For each iteration *k* = 1..*K*, we keep only the trajectories contained inside the square Γ_*k*,Δ*x*_ of size [(2*k* + 1) × (2*k* + 1)](Δ*x*)^2^ and centered at point *μ*_*k*–1_. The point *μ_k_* is the center of mass of points falling in the square Γ_*k*_ (eq. 14) and *μ*_0_ is the center of the initial high-density bin. The elliptic semi-axes *a_k_, b_k_* are computed as the *x*% confidence ellipse (eq. 17) from the covariance matrix *C_k_* (eq. (15)) and the angle *φ_k_* is the orientation of the ellipse (eq. 18). These parameters define the elliptic boundary of the well at iteration *k*

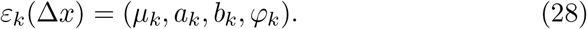

The field coefficient *A_k_* and diffusion coefficient *D_k_* are computed from eq. 26 and eq. 27 respectively, for the trajectories contained inside *ε_k_*. Specifically, we obtain *A_k_* from

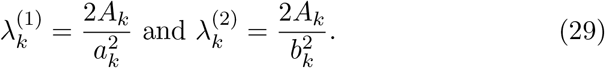

We repeat this procedure *K* times, with 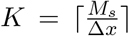 for the spatial parameter Δ*x* and the maximum region size *M_s_,* defined by the user.

##### Termination

This step consists in selecting the best iteration among *K*: we evaluate for each iteration *k* > 1 the likelihood score 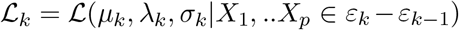 defined by eq. (24) but computed for sub-trajectories falling in the ring formed by the ellipses *ε*_*k*–1_ and *ε_k_*. The best iteration *k*\* is selected as the first local maximum of the curve 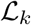.

#### 6.3.2 Multiscale Hybrid algorithm (MSHA)

We generalize the hybrid density-drift (FSHDD) algorithm defined above for a fixed spatial scale, by now varying the grid size Δ*x*, in the range Δ*x*_1_ < Δ*x*_2_ <.. < Δ*x_N_* selected by the user. The purpose of this MHSA is to select the optimal size Δ*x_i_*\* that minimizes the error

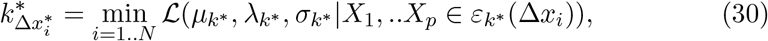

where 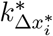 is the iteration that minimizes 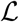 across all the spatial scales Δ*x_i_*, *i* = 1..*N*.

### 6.4 Density-Based Algorithm

The Density-Based Algorithm (DBA) uses the density of points estimated around the local density maximum of a high-density region. The algorithm uses the idea of level set of the Gaussian density distribution of points inside a potential well. We define the level set Γ_*α*_ with respect to a local maximum *M*\* as the ensemble of all trajectory points falling in bins with a density greater than *αM*\*:

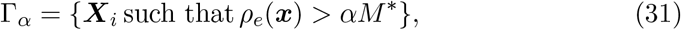

where *ρ_e_* is the empirical point density, estimated over the bins of a square grid (eq. 10) and *α* ∈ [0,1] is a density threshold. For the points 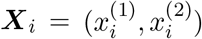 located in Γ_*α*_, the center ***μ*** of the distribution is approximated by the empirical estimators based on eq. 14 but restricted to the points in Γ_*α*_:

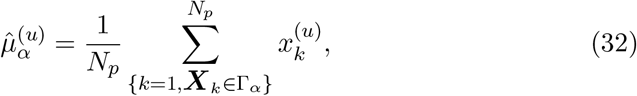

with *N_p_* the number of points in the ensemble Γ_*α*_ and *u* = 1..2. Similarly, we extend the estimator for the covariance matrix Σ (eq. 15) to

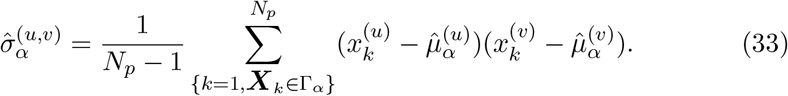

We now define the density-based algorithm:

#### Initiation

Search for high-density regions: the image is partitioned by a grid *G*_Δ*x*_ with square bins of size Δ*x* from which we compute the density map *ρ*Δ_*x*_(*x*) (eq. (10)). We then select the bins from *ρ*Δ_*x*_(*x*) with the highest *d*% density as possible regions containing a potential well.

#### Iterations

For each selected high-density bin, we initiated the well center 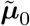 at the center of the bin. We then construct a refined grid centered at 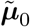 and with bin size Δ*x/p*, where *p* is chosen by the user (usually around 2 to 4). In this grid, we compute the centers ***μ***_0*α*_*k*__, for different values of *α*: *α*_1_ < … < *α_N_*, selected by the user. The refined center ***μ***_0_ is obtained as the center of mass of the centers ***μ***_0*α*_*k*__ for *k* = 1..*N*.

We then apply an iterative procedure that considers increasingly larger concentric annulus of center ***μ***_0_, width Δ_*r*_ and radius *r_k_* for *k* = 1..*K*. Where the number of iterations *K* is determined based on a minimal *r*_1_ = *r_min_* and maximal *r_K_* = *r_max_* distances defined by the user. For each iteration *k*, we compute the confidence ellipse *ε_k_* = (***μ***_0_, *a_k_*, *b_k_, φ_k_*) (see sub-section 6.2.4) obtained from the covariance matrix Σ_*k*_ (eq. 33) computed only for the points that fall in the annulus of radius *r_k_*. We then search for the iteration r* that maximizes the ratio 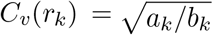 and use it to define the refined distance to the center

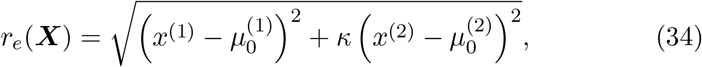

where *κ* = *C_v_*(*r*\*), that transforms an ellipse into a circle with the same center.

Finally, we compute the density of points *N_e_*(*r_k_*) falling in the annulus of radius *r_k_* based on the refined distance measure *r_e_*.

#### Termination

We select the first iteration *k*\* such that 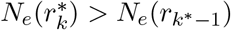: it is the first iteration where the derivative of the density with respect to the distance to the center, stops decreasing. This criteria is more stable on empirical data than searching for the minimum of the density (see figs. 3&4 panel B3 of [33]). The elliptic boundary of the well *ε*\* is centered at ***μ***_0_, has semi-axis lengths *a*, b\** given by *a*\* = *r_k_*\* and *b*\* = *κr_k_*\* and orientation *φ_k_*\*. We then use the ML estimator (eq. 27) to estimate the constant diffusion coefficient *D* inside *ε*\*. Finally, to compute *A*\* we use the diagonal form of the covariance matrix estimated (eq. 33) for all the points falling in *ε\**:

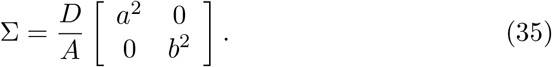

and estimate

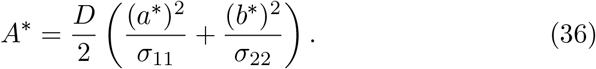

### 6.5 Drift-based algorithm

The drift based algorithm consists in using an error function in the space of the vector field to estimate the characteristics of a well.

#### Initiation

Search for high-density regions: the image is partitioned by a grid *G*_Δ*x*_ with square bins of size Δ*x* from which we compute the density map *ρ*Δ_*x*_(*x*) (eq. (10)). We then select the bins from *ρ*Δ_*x*_(*x*) with the highest *d*% density as possible regions containing a potential well.

#### Iterations

For each region selected in the initiation, we apply the following iterative procedure for *k* = 1..*K*:

a. We select only the sub-trajectories falling into a square *S_k_*(***μ***_*k*_, Δ*x*) centered at ***μ***_*k*–1_ and of size (2*k* + 1)Δ*x* × (2*k* + 1)Δ*x*. We take ***μ***_0_ to be the center of the high-density bin.
b. We estimate the elliptic well boundary *ε_k_* = [***μ***_*k*_, *a_k_, b_k_*, *φ_k_*] as the *X*% confidence ellipse (see sub-section 6.2.4) from the cloud of points in *S_k_*(*c_k_*, Δ). Where *X* is a parameter selected by the user (usually 90, 95 or 99).
c. We then compute a new grid *G*Δ_*x*_(***μ***_*k*_) centered at ***μ***_*k*_, that we use to estimate the local drift map (eq. 6) 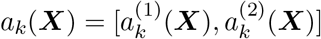 and estimate the attraction coefficient *A_k_* of the well using the least-square regression formula

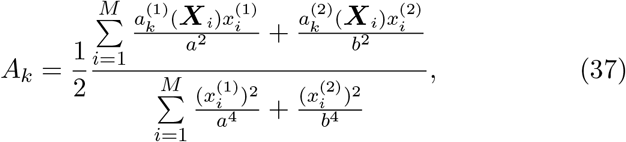

where 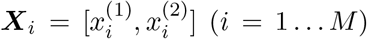 are the centers of the *M* bins from *G*_Δ*x*_ (***μ***_*k*_) that are contained inside the ellipse *ε_k_*.
d. Finally, we estimate the quality of the well (parabolic index) based on the residual least square error:

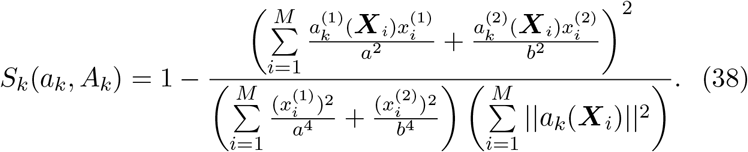

The index *S_k_* ∈ [0,1] is defined such that *S_k_* → 0 for a drift field generated by a parabolic potential well and *S_k_* → 1 for a random drift vector field as observed for diffusive motion.

The number of iterations is given by *N* = ⌊*w_max_*/Δ*x*⌋ where *w_max_* is the maximum size of the region to consider and is given by the user.

#### Termination

We select the iteration *k*\* that minimizes the parabolic index *S*: *k*\* = argmin_*k*=1…*K*_ *S_k_*(*a_k_, A_k_*). We estimate the diffusion coefficient inside the well using the local estimator (eq. 4) for all the displacements inside the ellipse *ε_k_*\*.

### 6.6 Sliding window analysis to study the stability of the wells over time

To determine the stability of the potential wells, we use a non-overlapping sliding window of 20s [14], to recover the ellipse characteristics, as shown on different examples in Supplementary Fig. 3A,C. When a well disappears in a given time window, but reappears latter, we kept the well for the entire period.

### 6.7 Reconstructing a graph for a network explored by SPTs

We describe here an algorithm to reconstruct a graph of a network explored by SPTs. This algorithm is based on the heterogeneous distribution of points caused by trajectories spending more time inside nodes than in tubules.

#### 6.7.1 Velocity based graph reconstruction algorithm (Vebgral) of a network explored by SPTs

The Vebgral algorithmic procedure to detect nodes (junctions) and inter-junction links (tubules) uses the large amount of recorded SPTs. The present algorithm first generates an ensemble of points from slow trajectory segments based on a maximum displacement length threshold *v_L_* and then uses the `dbscan` algorithm [36] to cluster these points based on their local density. The algorithm requires specification of two ensembles of parameters:

1. An ensemble of distances *R* = {*R_u_,u* = 1..*U*} (in *μ*m) defining the neighborhood radius around points.
2. An ensemble of counts *N* = {*N_v_, v* = 1..*V*} defining the numbers of points required in the neighborhood to form a cluster [36].

A pair of these two parameters (*R_u_, N_v_*) for any *u, v* define a local density 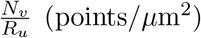 inside each cluster. The values of *R* and *N* depend on the local number of recorded trajectories and can vary inside the image. For each dataset, these values can be determined such that the computed clusters overlap with the structure of the organelle formed by the trajectories.

We now present the steps of the algorithm:

1. We form the ensemble of points belonging to low-velocity trajectory fragments 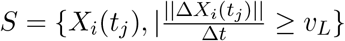.
2. We apply the `dbscan` procedure with parameters (*R_U_*, *N*_1_) to obtain a first ensemble of *K* clusters *c*_1_,…, *c_K_* from the points in *S*.
3. We then refine these initial clusters by searching for clusters possessing more than *N_max_* points. For any cluster *c_k_* possessing more than *N_max_* points:

a. We iteratively re-apply the `dbscan` algorithm with the more stringent parameter pair (*R_U_, N_v_*) for *u* = 2..*U* and *v* = (*V* – 1)..1 and replace the initial cluster with the resulting sub-cluster(s). We continue iterating over the generated sub-clusters until they all possess less than *N_max_* points.
4. We then approximate the boundary of each cluster either by its maximum volume ellipsoid or its convex hull polygon and assign each point discarded in step 1 to the cluster in which they fall if possible.
5. Finally, we merge any overlapping pair of clusters by computing the boundary of the combined ensemble of points (either elliptic or the convex hull) and we iterate this procedure until no more clusters overlap.

This first step allows to find the *K* nodes of the network. In the second step, we define tubules by constructing a connectivity matrix *C* of size *K* × *K* where *c_i,j_* (1 ≤ *i, j* ≤ *K*) is the number of trajectory displacements starting in node *i* and arriving in node *j*. Specifically, we increment *c_i,j_* for each data point *X_k_*(*t_l_*) (1 ≤ *k* ≤ *N_t_*, 0 ≤ l < *M_k_* – 1) in the following cases:

1. *X_k_*(*t_l_*) is located in node *i* and *X_k_*(*t*_*l*+1_) in node *j*
2. *X_k_*(*t_l_*) is located in node *i, X_k_*(*t*_*l*+1_) does not belong to any node and *X_k_*(*t*_*l*+2_) is located in node *j* (in this case 0 ≤ *l* < *M_k_* – 2).

### 6.8 Lysosome analysis

#### 6.8.1 Trajectories analysis for lysosome SPTs

To study the dynamics of lysosomes, we plotted the distribution of instantaneous velocities, computed from each trajectory displacement ***X***(*t* + Δ*t*) – ***X*** (*t*) by

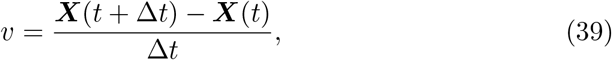

where Δ*t* = 1.5*s*. We approximate the distribution of instantaneous velocities using a two exponential model obtained by fitting 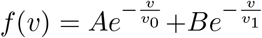 to the distribution using MATLAB’s `fit` toolbox. The density and diffusion maps were computed using the estimators 4 described above.

#### 6.8.2 Local high-density region analysis: ellipse approximation of the boundary

High-density regions are extracted from trajectories as follows: we construct a density map (eq. 10) based on a square grid with bin size Δ*x* = 480 nm. From this density map, we select only the 5% highest density bins and in case multiple such bins appear within a distance of two squares of each other, only the one with the highest value was kept. For each selected bin of center *c*, we computed a refined density map of size 5 × 5 squares, centered at *c* and with bin size Δ*x*′ = 200 nm. From this local map, we collected trajectory points falling into the bins that have a density > 80% of the maximal bin value [33] and use them to estimate the elliptic boundary of the region from the 95% confidence ellipse (see sub-section 6.2.4). Finally, when a pair of ellipses overlap, we replaced them by the ellipse computed over their combined ensemble of points and iterated this procedure until no more overlaps could be found.

#### 6.8.3 Transient confinement detection

To detect transient confinement periods along individual trajectories, we used the following procedure: for each point *Xt_j_* of a trajectory, we considered the ensemble of its successors *e_t_j,n__ = {*X*(t_j_*),…,*X*(*t*_*j*+*n*_)}, where initially *n* = *N_nh_* is set by the user. We then computed the center of mass *μ_t_j,n__* and checked that all the points *X*(*t_k_*) ∈ *e_t_j,n__* have a distance to the center of mass ||*X*(*t_j_*) – *μ_t_k,n__*|| < *R_nh_*, for a chosen distance threshold *R_nh_*(||.|| is the Euclidean norm). We then iterate the procedure, considering increasingly larger ensembles of successors *n* = *N_nh_, N_nh_* + 1, …*N_nh_* + *K* until either reaching the end of the recorded trajectory or when the next point ***X***(*t*_*j*+*n*+1_) do not fall into the neighborhood of the center of mass *μ_t_j,n__*. The confinement duration is then computed by considering the difference *t_*j+n*_* – *t_j_* in time between the two endpoints of the ensemble. Finally, the spring constant λ and diffusion coefficient *D* of the confinement is obtained by applying the Ornstein-Ulenbeck maximum likelihood estimators [48, 33], where the OU-process is given by

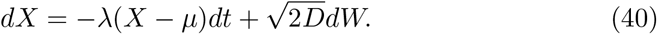

### 6.9 ImageJ plugin

The present method and algorithms are implemented into a imageJ plugin called “TrajectoryAnalysis”. The plugin allows to reconstruct the various maps (trajectories, density, drift, diffusion), detect potential wells and reconstruct the graph associated to trajectories. It allows to extract various statistics such as the distribution of diffusion coefficients or the energy and the size of potential wells.

## 7 Author contributions

PP and DH conceived and designed the research plan, computational methods and algorithms and wrote the manuscript. PP wrote the algorithms. M.H., E.A. J.H. and MH designed the experiments and collected superresolution microscopy data.

## 8 Acknowledgement

D. H.’s research has received funding from the European Research Council (ERC) under the European Unions Horizon 2020 research and innovation programme (grant agreement No 882673), Plan Cancer-INSERM Projet 19CS145-00 and ANR NEUC-0001.

## Notes

### Competing Interest Statement

The authors have declared no competing interest.

### Summary of Updates

we reduced the figure size and added the SI.

