## Supplementary methods for "SPtsAnalysis: a high-throughput super-resolution single particle trajectory analysis to reconstruct organelle dynamics and membrane re-organization"

November 7, 2021

The SI is organized as follows. First, we present the numerical validations of the reconstruction algorithm using a ground truth data set generated by an Ornstein-Ulhenbeck process. In the second part, we present complementary procedures to extract potential wells from calcium voltage channel trajectories in neurons.Third, we present further developments to reconstruct a graph map and possible potential wells that we apply to the dATL-mutant of the endoplasmic reticulum.

#### 1 Validation of the hybrid algorithm on synthetic trajectories

To validate the hybrid algorithm (section 1 Main text), we first define a ground-truth data set of trajectories. This ensemble is generated by numerical simulations of a stochastic equation considering a single potential well.

##### 1.1 Ground-truth data generated by a stochastic process and the associated numerical simulations

We define the ground-truth data as the ensemble of trajectories obtained by a truncated Ornstein-Uhlenbeck process, where the drift term is the gradient of a potential energy function given in eq. 12 from Main text. For a well of

*[1] Group of Data Modeling and Computational Biology, IBENS, Ecole Normale Supérieure,75005 Paris, France. [2] Research Group Functional Neurobiology at the Institute of Developmental Biology and Neurobiology, Johannes Gutenberg University Mainz, Mainz, Germany and [3] DAMPT, University Of Cambridge, DAMPT and Churchill College CB30DS, United Kingdom.* equally.

center $\boldsymbol{\mu}$ and boundary $\mathcal{B}$, the position $\boldsymbol{X}(t)$ of the process at time $t$ satisfies the stochastic equation

$$\dot{\boldsymbol{X}} = \begin{cases} -\lambda(\boldsymbol{X}(t) - \boldsymbol{\mu}) + \sqrt{2D}\dot{W} & X \in \mathcal{B} \\ \\ \sqrt{2D}\dot{W} & \text{otherwise.} \end{cases}, \tag{1}$$

where the diffusion coefficient is $D$ ($\mu m^2/s$) and the attraction coefficient is $\lambda$ ($s^{-1}$). We use the classical Euler's scheme to discretize the equation:

$$\boldsymbol{X}(t+\Delta t) = \begin{cases} -\lambda(\boldsymbol{X}(t) - \mu)\Delta t_{num} + \sqrt{2D\Delta t_{num}}N & X \in \mathcal{B} \\ \\ \boldsymbol{X}(t) + \sqrt{2D\Delta t_{num}}N & \text{otherwise.} \end{cases} \tag{2}$$

The random variable $N = [n1, n2]$ is Gaussian with $n1, n2 \sim \mathcal{N}(0,1)$ and $\Delta t$ is the elementary time step. Because experimental trajectories are obtained with a much coarser time step $\Delta t_{Exp} > 10$ ms, we first generated trajectories with a smaller time step $\Delta t_{num} = 1$ ms and then sub-sample the simulated trajectories, retaining one point every $\Delta t_{Exp}/\Delta t_{num}$. We generated the simulations with a diffusion coefficient $D = 0.05$ $\mu m^2/s$.

#### 1.2 Instantiation of the numerical simulations and estimation results

To simulate the ground truth trajectories for a well with boundary $\varepsilon = (\boldsymbol{\mu}, a, b, \varphi)$, we define a rectangular region $B$, for generating the initial trajectory points, centered at $\mu$ and of size $3a \times 3b$. Then, we generated trajectories uniformly distribution in the region $B$ but outside the well, until collecting 2000 displacements outside the well. After that, we generated trajectories uniformly distributed anywhere in the region $B$, until 1000 displacements have been collected inside the well. Each trajectory is composed of exactly 20 points, each separated by a time $\Delta t$ (fig. S1A).

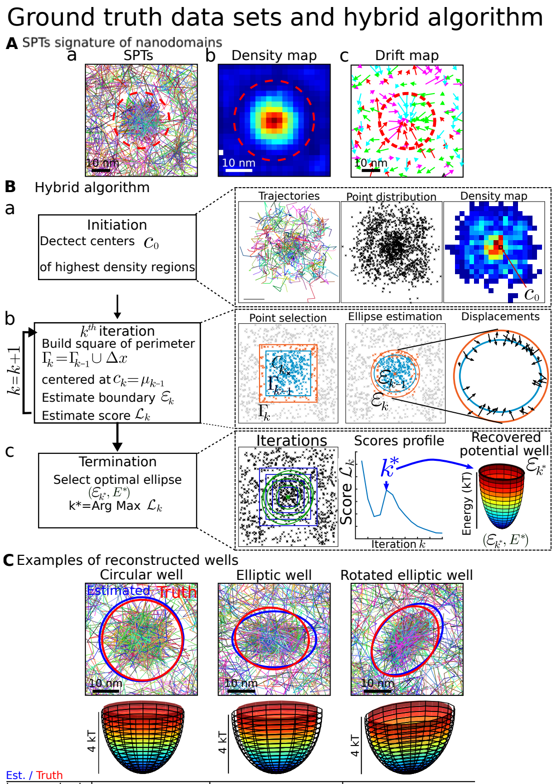


| Est. / Truth | Circular well | Elliptic well | Rotated elliptic well |
|---|---|---|---|
| large semi-axis (nm) | 145 / 150 | 132 / 150 | 139 / 150 |
| small semi-axis (nm) | 143 / 150 | 109 / 100 | 103 / 100 |
| Energy (kT) | 4.2 / 4.0 | 4.2 / 4.0 | 4.0 / 4.0 |

Figure S1: **Reconstruction algorithm of nanodomains from super-resolution SPTs. A.** a- Simulated trajectories attracted in a well of known boundary. b-density map c-drift field. **B.** Algorithm to reconstruct the center, the boundary and the energy of the nanodomain characterized as potential well. Initiation of the center $c_0$ with an ellipse $\gamma_0$ and the center of mass $\mu_0$. The error between the vector field and the reconstructed field is measured by the score $S_0$. The spatial resolution is fixed by $\Delta x$. k-iteration step: the domain is enlarged to $\Gamma_k$, the center is revaluated at point $c_k$ and the error $S_k$ is recomputed. In the termination step, the optimal value $k^*$ is selected that minimizes the error $S_{k^*}$, leading to the optimal center and ellipse $\gamma_{k^*}$, for which the energy is computed as the ratio $A/D$ of the field to the diffusion coefficient.

We generated different ensemble of parameters: considering a circular (of radius $r = 150$ nm) and an elliptic boundary (with semi-axes lengths $a = 150$ nm, $b = 100$ nm); For two typical values for the energy $E = A/D$ of $E = 4$ kT (low, $A = 0.2$) and $E = 6$ kT (high, $A = 0.3$), keeping the diffusion coefficient $D = 0.05$ $\mu m^2$/s constant; And finally setting the acquisition time $\Delta t = 20$ ms and $\Delta t = 50$ ms. For each condition, we generated 100 repeats of the simulations. We then used these datasets to test the capacity of our three algorithms (see Material and Method): hybrid algorithm (MLE), the algorithm based on the drift estimation(drift) and the one based on the density of points (Dens) to estimate from the trajectories the geometry of the well (semi-axes), the diffusion coefficient, the associated energy and the field coefficient $A$.

The results are shown in Tables 1.2 and 1.2 and we found that the hybrid algorithm (Fig. S1B) allows to recover the main parameters with a 10% error compared to the Drift and Density algorithms. Interestingly, the recovery of the main semi-axis depends on the energy $E$ of the well and the time step $\Delta t$ of the simulations. While the mean estimations of the semi-axes are satisfactory for an energy of $E = 4kT$, the estimated means are below the ground truth for the disk by less than 10% when applying the algorithm for $E = 6$ kT, independently of the time step. However, the mean value estimated for lower axis was above the true one, while the larger axis was sub-estimated for the elliptic case. Since the diffusion coefficient is well estimated, the possible errors found in the energy originates from the fluctuation in the estimation of $A$.

| | Algo | DT=20 ms | | DT=50 ms | |
|---|---|---|---|---|---|
| | | E=4 kT | E=6 kT | E=4 kT | E=6 kT |
| a (nm) | MLE | 152 ± 16 | 138 ± 21 | 158 ± 17 | 139 ± 26 |
| | Drift | 136 ± 12 | 135 ± 10 | 142 ± 12 | 142 ± 11 |
| | Dens | 145 ± 24 | 145 ± 11 | 153 ± 14 | 148 ± 8 |
| b (nm) | MLE | 145 ± 15 | 132 ± 20 | 152 ± 17 | 133 ± 24 |
| | Drift | 126 ± 12 | 126 ± 10 | 132 ± 12 | 134 ± 11 |
| | Dens | 139 ± 23 | 139 ± 13 | 149 ± 14 | 144 ± 9 |
| A ($\mu$m$^2$/s) | MLE | 0.23 ± 0.03 | 0.27 ± 0.06 | 0.24 ± 0.04 | 0.27 ± 0.08 |
| | Drift | 0.12 ± 0.03 | 0.17 ± 0.03 | 0.09 ± 0.02 | 0.13 ± 0.02 |
| | Dens | 0.16 ± 0.02 | 0.22 ± 0.02 | 0.19 ± 0.02 | 0.26 ± 0.02 |
| D ($\mu$m$^2$/s) | MLE | 0.050 ± 0.002 | 0.051 ± 0.002 | 0.050 ± 0.003 | 0.050 ± 0.004 |
| | Drift | 0.043 ± 0.001 | 0.041 ± 0.001 | 0.034 ± 0.001 | 0.029 ± 0.001 |
| | Dens | 0.049 ± 0.003 | 0.051 ± 0.002 | 0.050 ± 0.003 | 0.051 ± 0.003 |
| E (kT) | MLE | 4.51 ± 0.51 | 5.25 ± 1.06 | 4.75 ± 0.63 | 5.27 ± 1.49 |
| | Drift | 2.71 ± 0.57 | 4.07 ± 0.63 | 2.75 ± 0.46 | 4.41 ± 0.62 |
| | Dens | 3.35 ± 0.27 | 4.40 ± 0.29 | 3.80 ± 0.23 | 5.04 ± 0.31 |

Table 1: Recovered potential wells based on stochastic simulations for different sets of parameters and a circular well boundary ($a = b = 150$ nm), a field coefficient $A = 0.2\ \mu m^2/s$ (resp. $A = 0.3\ \mu m^2/s$) for $E = 4$ kT (resp. $E = 6$ kT) and a diffusion coefficient $D = 0.05\ \mu m^2/s$.

| | Algo | DT=20 ms | | DT=50 ms | |
|---|---|---|---|---|---|
| | | E=4 kT | E=6 kT | E=4 kT | E=6 kT |
| a (nm) | MLE | 134 ± 6 | 139 ± 6 | 149 ± 8 | 140 ± 30 |
| | Drift | 124 ± 11 | 125 ± 11 | 129 ± 9 | 127 ± 11 |
| | Dens | 136 ± 20 | 128 ± 14 | 142 ± 22 | 129 ± 16 |
| b (nm) | MLE | 110 ± 6 | 112 ± 6 | 122 ± 6 | 114 ± 27 |
| | Drift | 104 ± 8 | 100 ± 8 | 108 ± 7 | 104 ± 10 |
| | Dens | 116 ± 17 | 111 ± 14 | 123 ± 20 | 112 ± 15 |
| A ($\mu$m$^2$/s) | MLE | 0.23 ± 0.02 | 0.33 ± 0.03 | 0.26 ± 0.03 | 0.32 ± 0.07 |
| | Drift | 0.12 ± 0.02 | 0.16 ± 0.03 | 0.08 ± 0.01 | 0.10 ± 0.02 |
| | Dens | 0.17 ± 0.02 | 0.23 ± 0.02 | 0.18 ± 0.02 | 0.25 ± 0.04 |
| D ($\mu$m$^2$/s) | MLE | 0.051 ± 0.002 | 0.052 ± 0.003 | 0.050 ± 0.003 | 0.052 ± 0.008 |
| | Drift | 0.040 ± 0.002 | 0.036 ± 0.001 | 0.030 ± 0.001 | 0.023 ± 0.001 |
| | Dens | 0.050 ± 0.002 | 0.051 ± 0.003 | 0.048 ± 0.004 | 0.051 ± 0.007 |
| E (kT) | MLE | 4.49 ± 0.31 | 6.37 ± 0.52 | 5.12 ± 0.39 | 6.09 ± 0.83 |
| | Drift | 3.00 ± 0.44 | 4.34 ± 0.64 | 2.83 ± 0.40 | 4.29 ± 0.76 |
| | Dens | 3.50 ± 0.22 | 4.54 ± 0.30 | 3.75 ± 0.23 | 4.82 ± 0.36 |

Table 2: **Estimation of the potential well characteristics** using the hybrid algorithm and comparison with the Density and the Drift algorithms for an elliptic well. The parameters are the same as for Table. 1.2 except that we use an elliptic boundary with $a = 150$ nm and $b = 100$ nm.

### 2 SPTs analysis

#### 2.1 Increased accuracy of map reconstruction with cosine-window estimators

We compare here the diffusion map computed with the classical super-resolution estimators (Eq. 7 Main text) against the estimators based on the cosines filter (Eq. 9 Main text). The results are shown in Fig. S2, where the cosines estimators give a much higher spatial resolution, as it is based on two scales: a radius $r = 0.2\ \mu m$ for the moving disk, and a grid bin width $\Delta x = 0.05\ \mu m$. In this procedure, we keep the bins that contain at least 5 displacements.

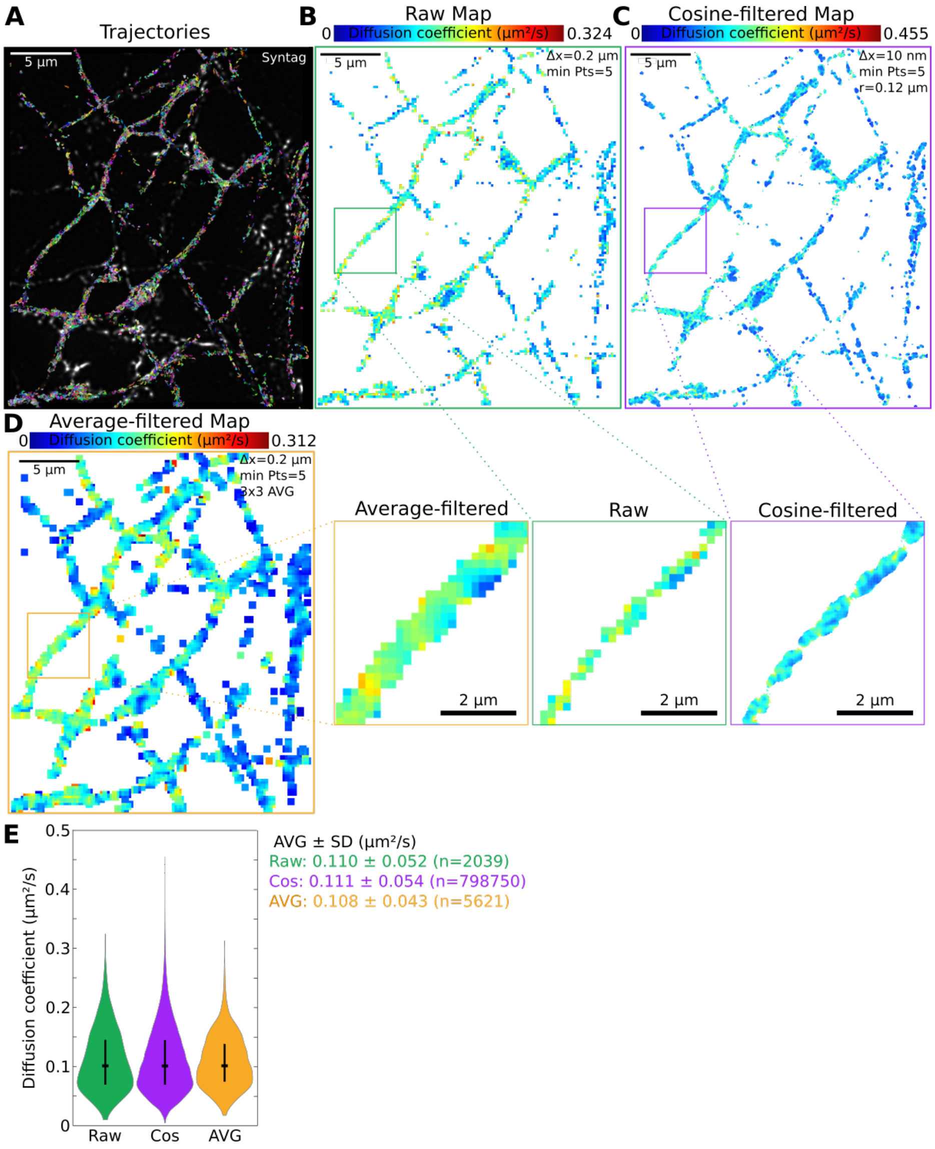


Figure S2: Comparing the diffusion map for CaV SPTs using three estimators. **A** CaV SPTs. **B** Diffusion map computed from the classical estimator (Eq. 7 main text). **C** Diffusion map computed with the cosine-estimator (Eq. 9 Main text). The cosine estimator is computed over a sliding window with a moving disk of radius $r = 0.2\ \mu m$ and the grid unit length is $\Delta x = 0.05\ \mu m$. **D**Map computed using a medium filtered (low pass using the mean of the 9 neighbors). Comparison of average-filtered vs single bin map and cosines-filtered map. The average-filtered diffusion map is computed from the diffusion map by replacing the value for each bin by the average value computed only over the bins possessing a diffusion coefficient in a 3x3 region around the original bin. **E** Violin plots of the diffusion coefficients computed in each bin.

#### 2.2 Time lapse analysis of SPT trajectories

We define the time lapse analysis as a sequence of possibly overlapping time windows $W_1, \ldots, W_N$ of duration $\Delta W$. We then split the ensemble of trajectories into these time windows and independently analyze them. A schematic example is presented Fig. S3, where we show a 50% overlap between windows. We applied this procedure when mentioned in the text.

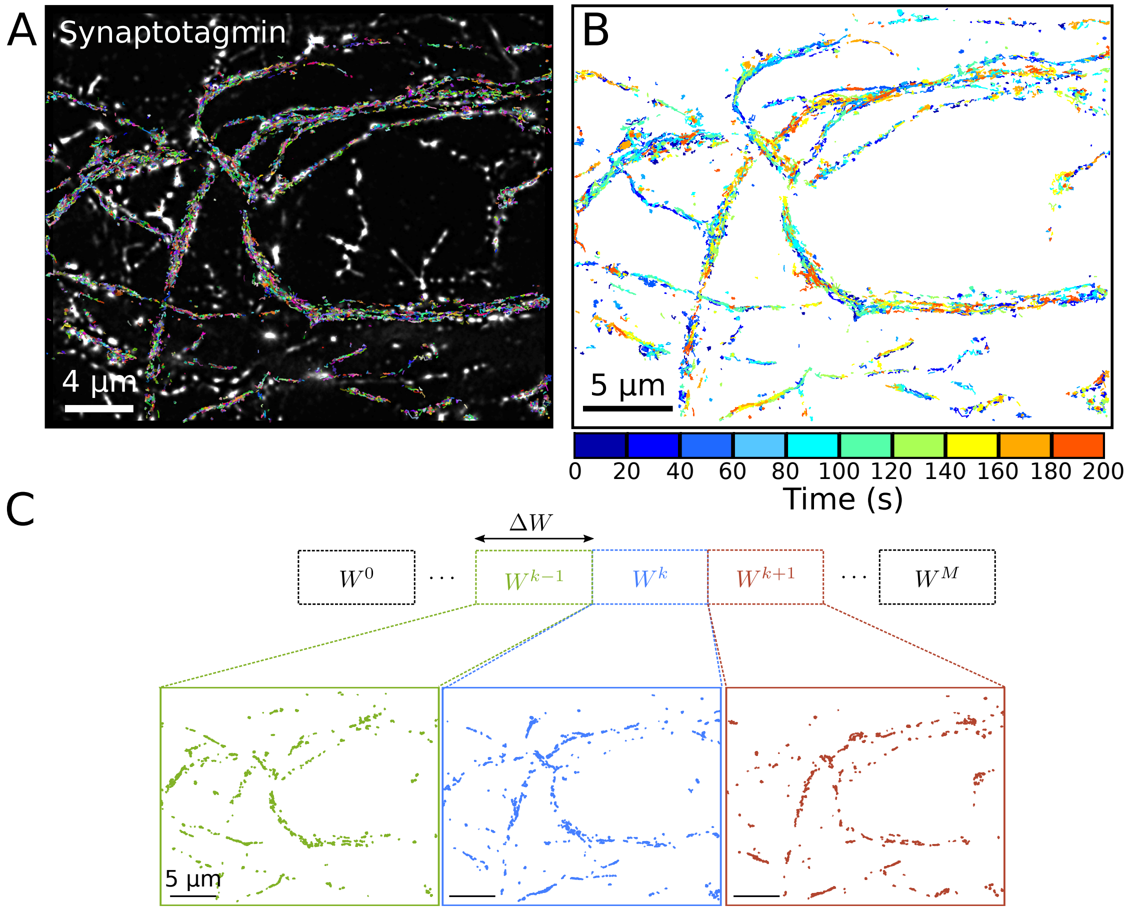


Figure S3: **Time Lapse analysis based on a sliding window A.** Example of a single-particle trajectory dataset plotted over the entire time acquisition. **B.** Time-splitting of the dataset into 20 s time windows with 50% overlap. Trajectories are colored by the time window where they appear. **C.** Time-splitting principle: the entire duration spanned by the dataset is divided into windows of duration $\Delta W = 20$ s with an adjustable overlap duration chosen here to be 10 s. A trajectory belonging to two consecutive time windows will serve for the statistical estimations for each period.

Table 2.2 summarizes the geometrical properties of the wells found in CaV variants. Here $\tau_e$ is the mean residence time of channels inside a well

| Exp. | a (nm) | b (nm) | D ($\mu m^2$/s) | A ($\mu m^2$/s) | E (kT) | $\tau_e$ (ms) | n |
|---|---|---|---|---|---|---|---|
| Cav2.1 Δ47 | 143 ± 51 | 104 ± 33 | 0.091 ± 0.052 | 0.345 ± 0.209 | 3.9 ± 1.1 | 174 ± 134 | 1587 |
| Cav2.1+47 | 145 ± 52 | 103 ± 33 | 0.087 ± 0.045 | 0.337 ± 0.189 | 3.9 ± 1.1 | 181 ± 141 | 1713 |
| Cav | 100 ± 47 | 73 ± 31 | 0.069 ± 0.051 | 0.224 ± 0.184 | 3.3 ± 1.2 | 94 ± 92 | 1877 |

Table 3: **Wells parameters extracted from CaV SPTs**

(see Main Text).
Table 2.2 summarizes the long-time stability of the wells.

| Exp. | Stability (s) | n |
|---|---|---|
| Cav2.1Δ47 | 56.2 ± 25.5 | 1105 |
| Cav2.1+47 | 57.4 ± 25.3 | 1167 |
| Cav | 46.6 ± 15.7 | 1574 |

Table 4: **Wells stability for CaV recorded for hippocampal neurons.**

#### 2.3 Tables associated to CaV SPTs and GFP-binding intrabodies CaV

| Exp. | Ratio of trajs. in wells | n |
|---|---|---|
| Cav2.1Δ47 | 0.418 ± 0.094 | 19 |
| Cav2.1+47 | 0.427 ± 0.129 | 19 |
| Cav | 0.328 ± 0.081 | 11 |

Table 5: **Ratio of trajectories appearing inside wells.** Results are given as AVG ± SD.

Fig.S4 shows CaV nanodomain automatically identified during a time lapse analysis.

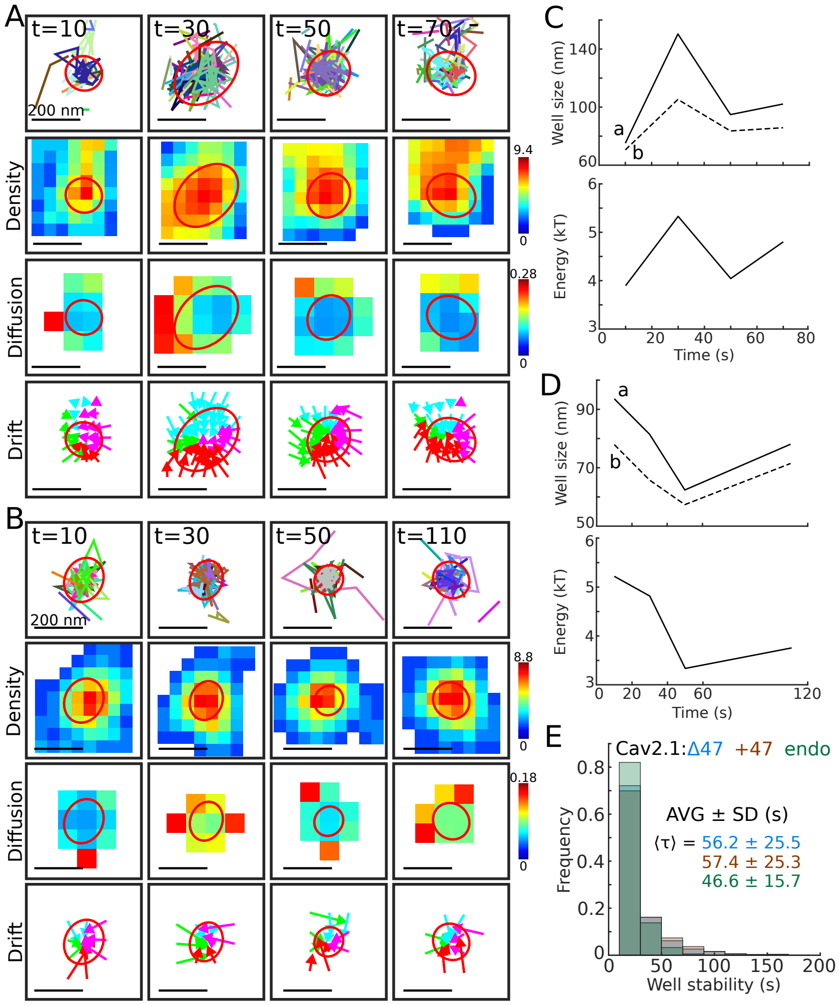


Figure S4: **2 examples of CaV nanodomain automatically identified during a time lapse analysis.** **A.** Example 1: high density region identified as potential well in successive 20 s time windows. From top to bottom: individual trajectories, associated density, diffusion and drift maps. **B.** A second example of a potential well, similar to A. **C-D.** Temporal evolution of size and energy characteristics associated to the well presented in A and B respectively. **E.** Population characteristics for three experimental conditions: CaV2.1-47, CaV2.1+47 or CaV2.1 showing the mean life time of potential wells extracted by an exponential fit.

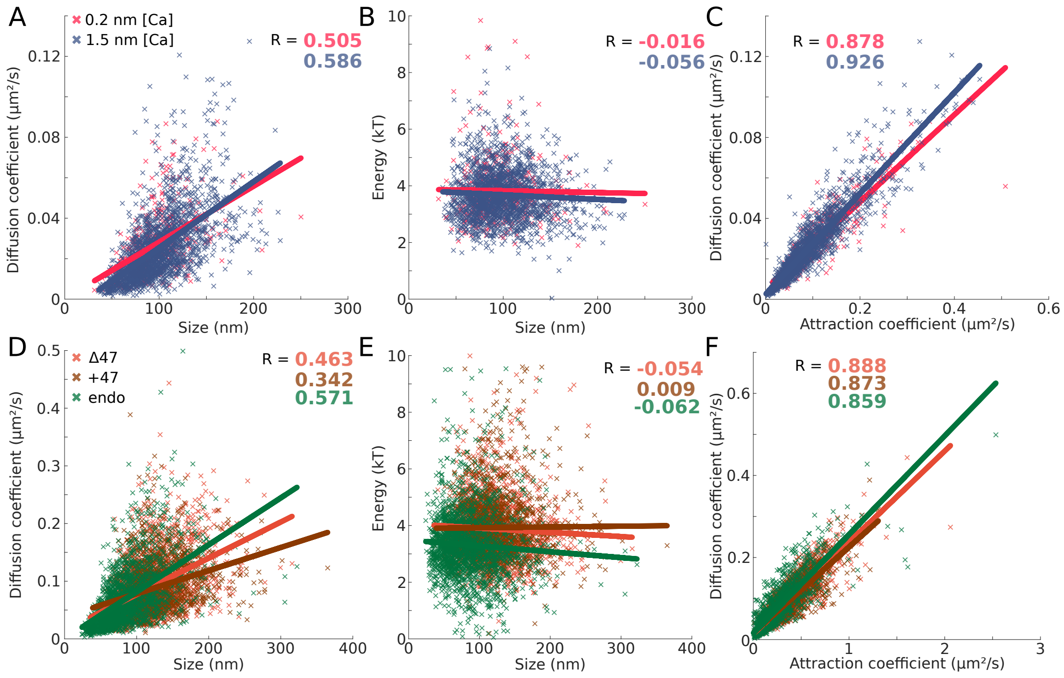


Figure S5: **Correlation statistics obtained inside CaV wells revealed by super-resolution SPTs. A.** Diffusion vs size (high correlation) **B.** energy vs size (no correlation). **C.** Diffusion vs size, positive regression $a_{+47} = 0.46$ $a_{-47} = 0.34$, $a_{endo} = 0.51$.**D.** Energy vs size (no correlation).

### 3 Biophysical characterization of ER nodes as potential wells to retain material

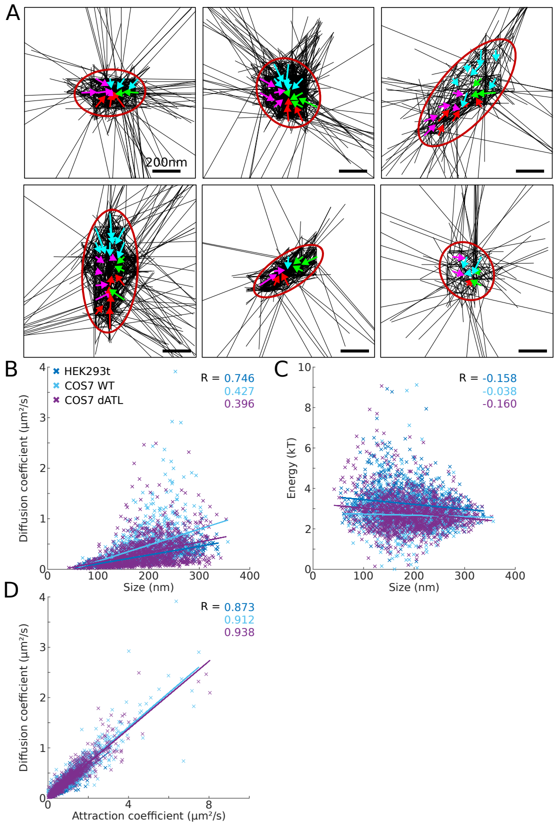


Figure S6: **ER nodes are revealed by super-resolution SPTs and characterized as potential wells. A.** 6 examples of converging arrows in potential wells associated to ER nodes. **B.** Distribution of the energy of the wells vs the size and the correlation coefficients: $C_{HEK-297} = 0.74$ $C_{Cos7-WT} = 0.42$, $C_{Cos7-dATL} = 0.39$. **C.** Energy vs size (no correlation), see also table 3.

| Exp. | a (nm) | b (nm) | D ($\mu$m$^2$/s) | A ($\mu$m$^2$/s) | E (kT) | $\tau_e$ (ms) | n |
|---|---|---|---|---|---|---|---|
| HEK293t | 219 ± 71 | 155 ± 56 | 0.252 ± 0.136 | 0.787 ± 0.401 | 3.3 ± 0.9 | 101 ± 64 | 1057 |
| COS7 WT | 230 ± 68 | 167 ± 45 | 0.461 ± 0.369 | 1.227 ± 0.984 | 2.7 ± 0.8 | 63 ± 46 | 859 |
| COS7 dATL | 225 ± 70 | 163 ± 54 | 0.321 ± 0.292 | 0.868 ± 0.814 | 2.8 ± 0.9 | 106 ± 89 | 1250 |

Table 6: **Recovered wells parameters for ER datasets for HEK293t,COS7 WT and COS7 dATL**.

##### 3.1 Dispersion index to estimate how trajectories explore locally the ER nodes

To investigate how trajectories disperse when exiting an ER node, we synchronize them on their last point spent in node $n$. Trajectories are described as $X_i = \{X_i(t_j)\}$ where $j = 1 \dots M_i$ is the number of points in trajectory $X_i$. For each trajectory $X_i$ passing through the node $n$, we define its synchronization time $t_i^*$ to be its last point inside $n$: $X_i(t_i^*) \in n$ and $X_i(t_{i^*+1}) \notin n$. The dispersion index is computed for each node $n$ on the ensemble of synchronized trajectories:

$$I_n(k) = \frac{1}{N_{n,k}} \sum_{i,j}^{i>j} ||X_i(t_{i^*+k}) - X_j(t_{j^*+k})||, \tag{3}$$

where $N_{n,k}$ is the number of unique pairs that can be formed from the remaining synchronised trajectories $k > 0$ frames after exiting node $n$ and $||.||$ is the Euclidean norm.

#### 4 Dynamical properties under dATL mutation for local trajectory exploration

Analysis of SPTs obtained from lumen in dATL mutant COS-7 cells reveal a change in the network organization (Fig. S7A) characterized by an extended rectilinear motion, suggesting that trajectories are now restricted in tubules that are of $\approx 7.5$ $\mu m$ in length.

##### 4.1 Long tubule detections from SPTs

We use the following method to detect long tubules from individual trajectories recorded inside the ER. This method is based on the assumption that these trajectories should be very elongated. Considering an ensemble

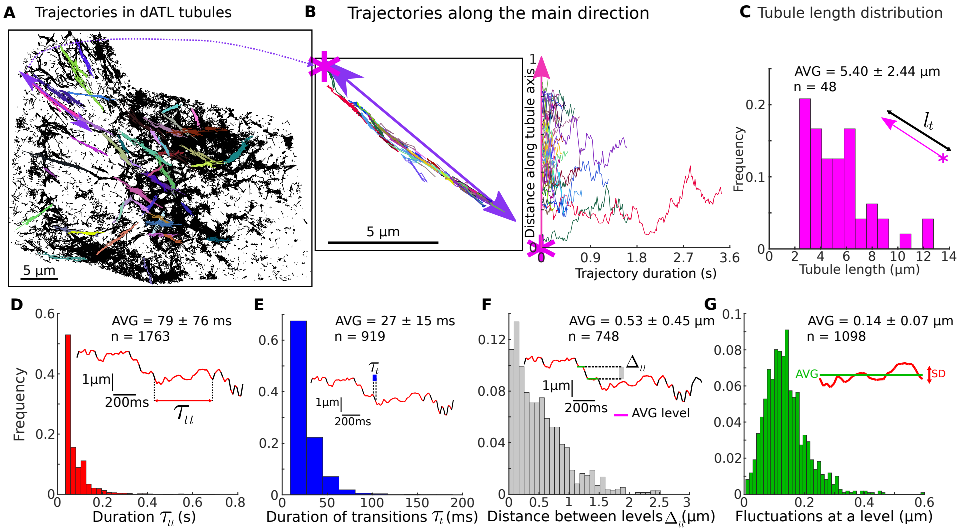


Figure S7: **Trajectory dynamics in long tubules from dATL cos7 cells.** **A** SPTs (black) recorded in the ER of a cos7 dATL cell with a detected long-tubules (colored). **B** Magnification of a single detected long tubule. **C** Distance vs time along the tubule traversed by trajectories. **C** Length distribution for long tubules presented in A. **D-G** Decomposition of the projected motion of trajectories in long tubules showing stalling and transition between different levels. **D** Duration distribution at a given level, **E** Distribution of transitions between two successive levels. **F** Distribution of distances between two successive levels. **G** Fluctuation amplitude of the positions inside a level.

of $N$ trajectories $\{X_i, i = 1, .., N\}$ such that $X_i = \{X_i(t_j), j = 1, .., M_i\}$, the algorithm is as follows:

1. We remove trajectories with less that $th_{traj\_size}$ points.

2. To estimate the amount of elongation of trajectories, we fit an ellipse $\varepsilon_i = (\boldsymbol{\mu}_i, a_i, b_i, \varphi_i)$, where $\boldsymbol{\mu}$ is the ellipse center, $a, b$ the large (resp. small) semi-axes lengths and $\varphi$ the orientation, around each trajectory $X_i$, $i = 1..N$ using the minimum volume ellipsoid method.

3. We keep trajectories $X_i$ such that the semi-axes ratio $th_{traj\_rat\_min} < \frac{a_i}{b_i} \leq th_{traj\_rat\_max}$.

4. We then compute the bounding square $s_i$ of the ellipse $\varepsilon_i$ of width $2a_i$, length $2b_i$, orientation $\varphi_i$ and center $\boldsymbol{\mu}_i$.

5. We then seek to form groups of overlapping trajectories with similar orientations. Considering the set of already processed trajectories $P$, we form a new group of trajectories $G$ as follows:

   (a) Start with the next unprocessed trajectory $i$, add it to $P$ and $G$: $P = P \cup \{i\}$, $G = G \cup \{i\}$.

   (b) While $G$ has changed since the last iteration, we do the following update:

      i. For each trajectory $j \in G$:

         A. Search any unprocessed trajectory $k \notin P$

         B. if the area of the overlap between the bounding square of the two trajectories $A_{j,k} = \text{area}(s_j \cap s_k)$ such that $A_{j,k} > th_{area\_rat} \times \min(\text{area}(s_j), \text{area}(s_k))$ and the angle between the two squares is $< th_{sq\_angle}$, then add $k$ to $G$ and $P$: $G = G \cup \{k\}$, $P = P \cup \{k\}$.

   (c) The resulting group is composed of trajectories with similar orientations that approximates well the observed long tubules.

6. Finally, we filter out groups with $< th_{group\_size}$ trajectories, and similarly to trajectories, we fit an ellipse $\varepsilon_g = (\boldsymbol{\mu}_g, a_g, b_g, \varphi_g)$ around all the trajectories in the group using the minimum volume ellipsoid method and filter out groups for which the ratio of semi-axes lengths $r_g = \frac{a_g}{b_g} < th_{group\_rat}$.

The parameters used for the above method are given in Table 4.1.

| Parameter | Value |
| --- | --- |
| $th_{traj\_size}$ | 5 |
| $th_{traj\_rat\_min}$ | 4 |
| $th_{traj\_rat\_max}$ | 40 |
| $th_{area_rat}$ | 0.2 |
| $th_{sq\_angle}$ | $\frac{\pi}{12}$ |
| $th_{group\_size}$ | 5 |
| $th_{group\_rat}$ | 11 |

Table 7: **Parameters used for the long tubule detection algorithm.**
